# DNA Origami Nanomechanical Amplifiers for Resolving Single-Molecule Binding Events

**DOI:** 10.64898/2026.08.11.743830

**Authors:** Rui Yee Loke, Lennart J.K. Weiß, Enzo Kopperger, Friedrich C. Simmel

## Abstract

Mechanical amplification of minute length changes enables precise measurements across many orders of magnitude, from macroscopic metrology to optical instrumentation. Extending this principle to molecular systems could provide a route to monitoring nanoscale structural changes without relying on analyte labeling or fluorescence-based distance measurements. Here we present a DNA origami nanomechanical amplifier that converts subnanometre-scale molecular conformational changes into amplified mechanical displacements that can be tracked in real time at the single-molecule level. The platform resolves geometric changes associated with DNA hybridization, secondary-structure formation, DNA strand-exchange dynamics, and ligand-induced aptamer folding, enabling quantitative analysis of molecular kinetics and direct observation of transient intermediates and heterogeneous conformational ensembles. By translating molecular recognition events into mechanically amplified signals, our approach establishes a general framework for monitoring binding-coupled conformational dynamics and extends the scope of single-molecule measurements beyond conventional optical readouts.

## Introduction

Single-molecule measurements provide access to molecular heterogeneity and dynamics that remain obscured in ensemble experiments, enabling direct observation of binding events and conformational changes at the level of individual molecules [1, 2]. Such capabilities are central to both fundamental studies of biomolecular processes [3, 4] and the development of sensitive biosensing platforms [5].

Established single-molecule force spectroscopy techniques, including atomic force microscopy and optical or magnetic tweezers, have yielded detailed insights into processes such as DNA mechanics [6–10], transcription [11, 12], and chromatin remodeling [13, 14]. However, these approaches rely on complex instrumentation and are inherently difficult to scale or parallelize. High-throughput alternatives based on fluorescence [15, 16] or nanopores [17, 18] offer increased parallelism and speed, but direct and quantitative determination of nanometre-scale distance changes remains challenging. For example, while single-molecule FRET [15, 19] provides access to distances in the nanometre range, extracting absolute and time-resolved distance changes in a quantitative manner is often nontrivial due to calibration uncertainties, fluorophore orientation effects, photophysical variability, and its limited dynamic range.

DNA origami provides a complementary route by enabling the construction of nanoscale devices with precisely defined geometry and programmable, site-specific functionalization. In particular, dynamic DNA origami structures can transduce molecular interactions into mechanical motion, offering a direct physical readout of binding-induced conformational changes [20, 21].

Hinge-like DNA origami devices exemplify this concept by functioning as molecular calipers: binding events near the hinge induce changes in the opening angle, which are geometrically amplified by extended structural units [22, 23]. This mechanical amplification converts nanometre-scale structural changes into a more readily measurable observable, enabling distance measurements using electron microscopy or fluorescence-based readouts. However, existing implementations have largely been limited to static measurements and do not capture the kinetics of molecular interactions [24], restricting their ability to resolve weak or transient binding events and subtle conformational transitions.

Here, we introduce a dynamic DNA origami device that couples target binding to mechanically amplified motion through an extended lever arm, enabling time-resolved single-molecule measurements of molecular interactions (Fig. 1a). By converting sub-nanometre conformational changes into an amplified, geometrically defined signal, our approach provides quantitative access to both the magnitude and kinetics of molecular interactions in a scalable single-molecule format.

**Fig. 1:**
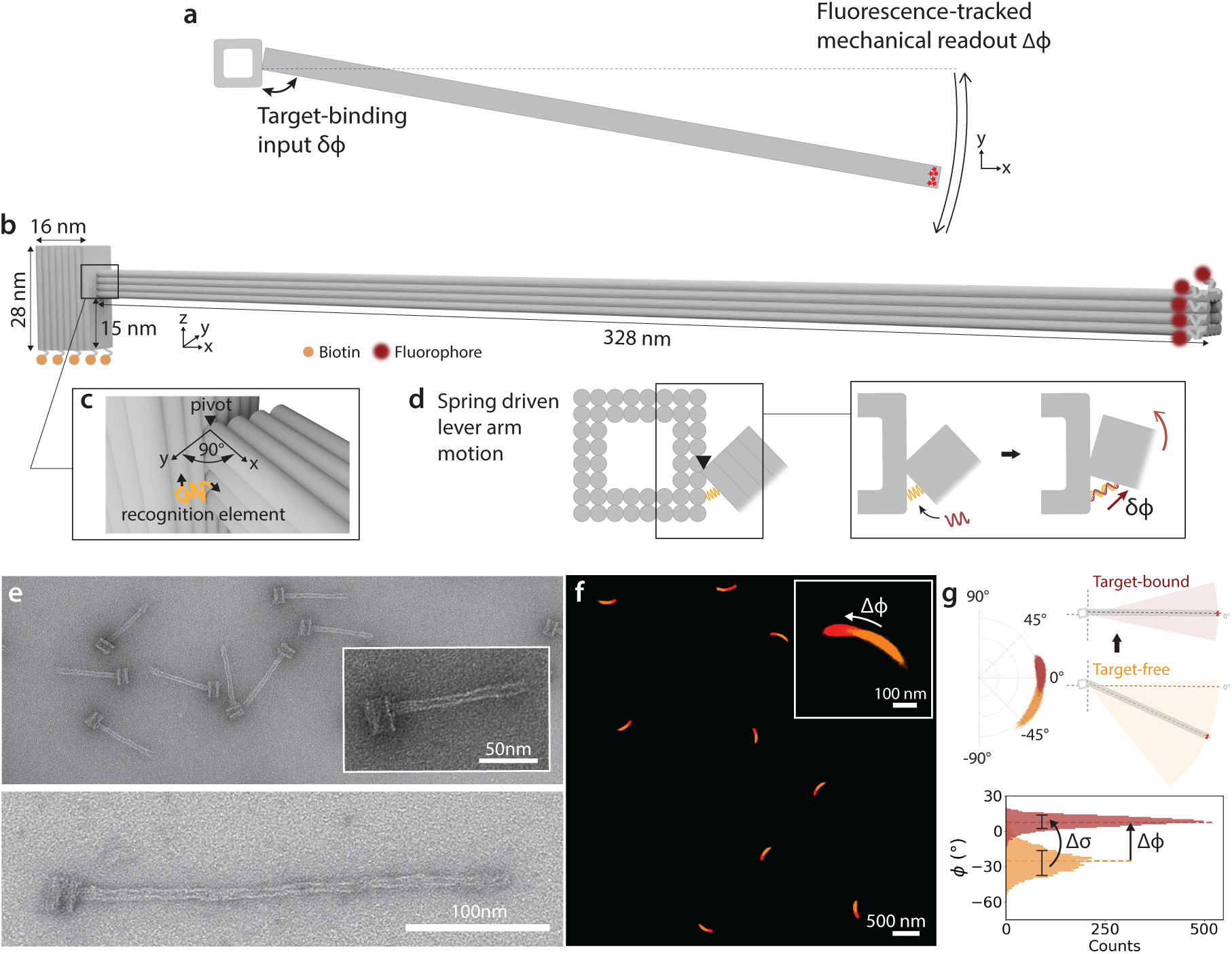
Nanomechanical amplifier design. **a**, The nanodevice leverages a lever arm mechanism, in which sub-nanometre conformational changes (*δϕ*) induced by target binding are amplified into optically resolvable mechanical readout (Δ*ϕ*). **b,** The approximately 300 nm-long lever arm is connected to a supporting base via four double-stranded crossovers that form a compliant hinge. The supporting base is immobilized on the surface via biotin–NeutrAvidin coupling, thereby elevating the lever arm by *≈*15 nm to minimize nonspecific interactions. **c,** The hinge serves as a pivot for a *≈* 90*^◦^*planar-motion of the lever arm. Arrows indicate the locations at which the molecular recognition element is integrated into the lever arm and the base. **d,** The recognition element tethered between the lever arm and the base exhibits a spring-like compliance in response to target binding. Binding-induced spring deformation is converted into lever arm rotation about the hinge, which is amplified by the lever arm into real-time displacement at the distal tip. **e,** TEM micrographs of the monomer platform (top) and dimer assembly with lever arm extension (bottom). **f,** TIRFM localization heatmaps overlay showing lever arm motion before (orange) and after 21 nt target strand binding (red), with an enlarged view displaying the localization cluster corresponding to a single platform (angular positions shown in g). **g,** Re-referenced angular positions (top left), with top-view schematics illustrating lever arm motion in target-bound and target-free state (top right) and corresponding angular histograms (bottom). The binding-induced lever arm displacement Δ*ϕ* is determined from the difference between the mean angular positions in the target-bound 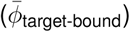 and target-free 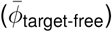 states. Target binding also alters the angular fluctuations, quantified by the change in standard deviation Δ*σ*.

We demonstrate that our setup can resolve sub-nanometre length differences by probing DNA hybridization of duplexes with base-pair (bp) resolution. We further show the device enables time-resolved monitoring of DNA hybridization between probes and targets, allowing detection of single-base mismatches and real-time observation of strand-exchange dynamics at the single-molecule level. Finally, our nanomechanical amplifier enables monitoring of aptamer folding and target binding at the single-molecule level, capturing the associated geometric length changes upon binding and resolving structural heterogeneity within the ensemble.

### DNA origami nanomechanical amplifiers

Our DNA origami–based nanomechanical sensing platform comprises a lever arm connected to an immobilized supporting base via four double-stranded linkers that form a compliant hinge (Supplementary Fig. S1), restricting motion to planar rotation over approximately 90*^◦^* (Fig. 1b,c). In previous studies of DNA origami rotors [25], as well as earlier versions of the geometric amplifier, we found that interactions between the lever arm and the substrate give rise to non-negligible unspecific interactions that interfere with the detection of target analyte interactions occurring on similar energy and time scales. Elevating the lever arm by approximately 15 nm above the substrate effectively overcomes this limitation.

The molecular recognition element is tethered at one end to the lever arm and at the other end to the base, generating a spring-like response to target binding (Fig. 1d). Contracting or stretching upon binding drives rotation of the lever arm around the hinge, which is geometrically amplified by the *≈*300 nm-long lever (see Supplementary Fig. S2) into a measurable displacement at the distal tip. The distal tip is fluorescently labeled and tracked with single-molecule precision using total internal reflection fluorescence microscopy (TIRFM). Transmission electron microscopy (TEM) confirms the structural integrity of both the monomeric platform and the dimeric assembly featuring an extended lever arm (Fig.1e). The supporting base is immobilized on the coverslip via biotin–NeutrAvidin coupling, positioning the lever arm above the surface to minimize nonspecific interactions during TIRFM measurements (Supplementary Fig. S4a). After immobilization, the sensor platforms adopt random orientations (Fig.1f). We therefore aligned the angular positions of all tracked fluorophore localizations to a platform-specific reference angle, enabling a standardized analysis across structures. From the angular distributions of each particle, we extracted changes in the mean angular position Δ*ϕ* and standard deviation Δ*σ* (Fig.1g). In a field of view of approximately 95 *×* 65 *µ*m^2^, 150-500 particles were selected, while excluding those with overlapping localizations or aberrant target-free localization patterns (Supplementary Fig. S3). To characterize the dynamics of the detection system, we analyzed the lever arm motion in the absence of a target, and found that the resulting angular fluctuations are well described by a Gaussian distribution, consistent with a spring-like system governed by a harmonic potential (Supplementary Fig. S5a). Notably, the nanomechanical amplifier can also be utilized as a transducer for steric hindrance events (Supplementary Figs. S6 and S7).

### Base pair-resolved mechanical readout

To probe the sensitivity of our system to small molecular length changes, we quantified the nanomechanical response induced by binding of DNA strands of lengths ranging from 7 to 21 nt in two-base increments (*≈*0.68 nm per step) to the 21 nt single-stranded DNA recognition site (Fig. 2a), with base increments progressing toward the 3’-end of the recognition strand tethered to the base (Fig. 2b). Measurements were performed at a target strand concentration of 200 nM (i.e., in excess over the recognition strands) following 5 min incubation in each round of independent measurements. Formation of a more rigid double-helical structure led to an anticlockwise lever arm displacement (Δ*ϕ >* 0) and reduced angular fluctuations (Δ*σ*; Fig. 2c-e). Δ*ϕ* values were extracted for individual platforms, and the median response across the ensemble (n=77) was determined for each target length to obtain a calibration curve for base-pair formation (Fig. 2d). Given the weak nonlinearity of the response, we approximated the response as linear, corresponding to an angular displacement of *≈* 2.5*^◦^* per two base pairs over the measured range. For a 300 nm lever arm, this angular change corresponds to a tip displacement of 13 nm (*s* = *rθ*, where *r* is the lever-arm length and *θ* is the angular displacement in radians). Because the formation of two base pairs produces a conformational change of only 0.68 nm, the nanomechanical amplifier provides an approximately 20-fold geometric amplification.

**Fig. 2:**
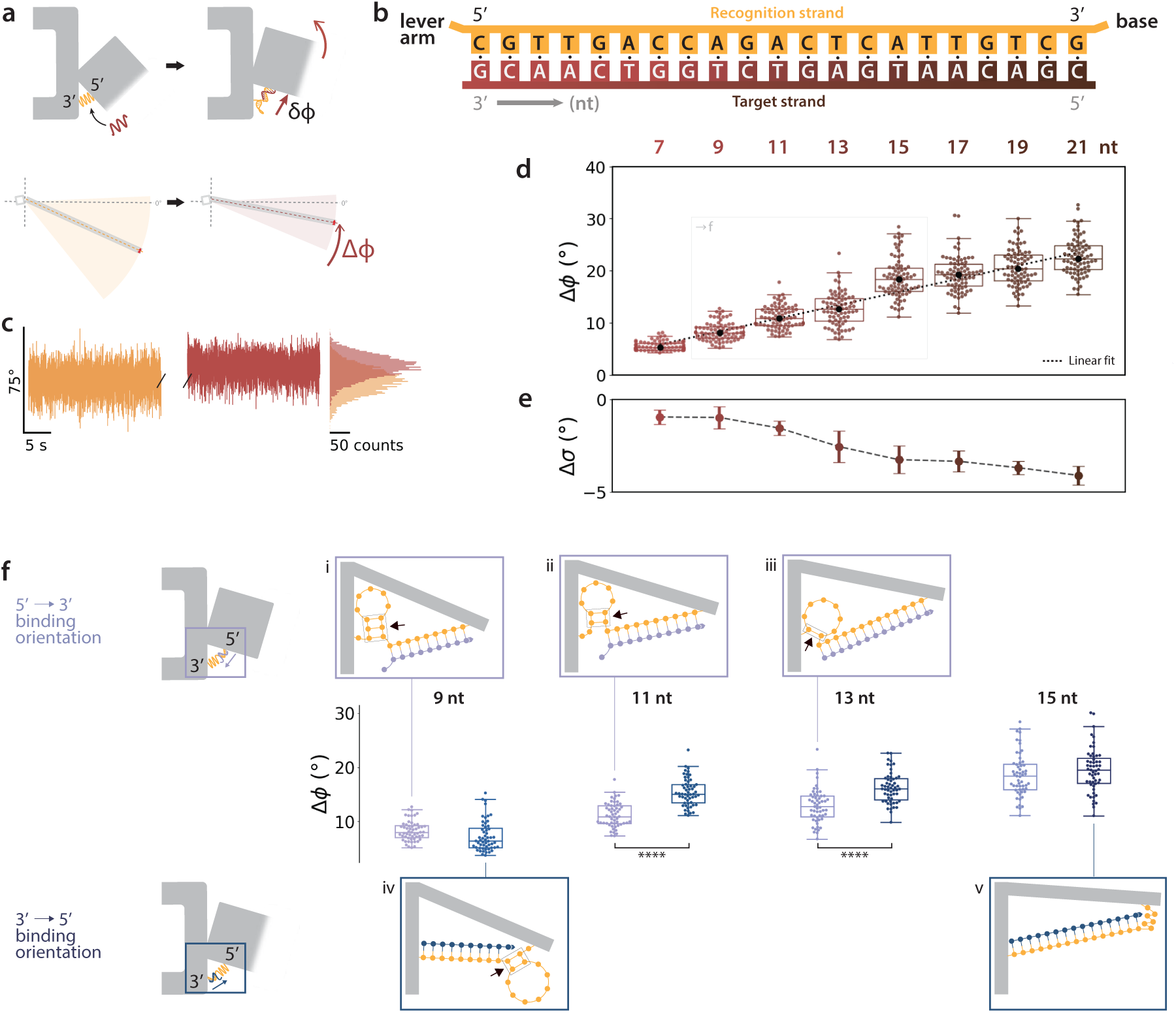
Angular response resolved at the base-pair level. **a**, Top-view schematic of the platform, illustrating target-strand hybridization to the recognition strand, resulting in an anticlockwise displacement of the lever arm (Δ*ϕ >* 0). **b,** Target strand lengths were increased in two-base increments toward the 3’-end of the recognition strand tethered to the base, ranging from 7 to 21 nt in each round of independent measurements. **c,** Representative single-lever arm time traces before (orange) and after (red) binding of a 9 nt target strand with corresponding angular histograms. **d,** Δ*ϕ* as function of the target length (n=77). Selected points (9–15 nt) are further analyzed in **f**. **e,** Binding-induced changes in lever arm fluctuations (Δ*σ*). **f,** Pairwise comparison of Δ*ϕ* for the same target length with different loose-end geometries in the recognition-target duplex (n=55). Statistical significance was assessed with two-sided Welch’s t-test with *α* = 0.05 Holm adjustment; \*\*\*\**P <* 0.0001. Schematics illustrate loose-end hairpin conformations of recognition-target duplex subject to 5’*→*3’ or 3’*→*5’ binding orientations for 9, 11, 13 nt **(i-iv)**. Unspecified duplexes have loose ends without hairpins, as shown in the illustration for 15 nt **(v)**. Schematics are conceptual; the corresponding NUPACK-predicted equilibrium conformations are shown in Supplementary Fig. S8.

Binding of progressively longer targets also confines the lever arm more tightly around its equilibrium position (Fig. 2e). This effect was quantified by the change in standard deviation Δ*σ*, which decreased with duplex length, from *−*1.0*^◦^* for a 7 bp duplex to *−*4.1*^◦^* for a 21 bp duplex. In our experiments, target strands ranging from 7 to 21 nt bind to the same 21 nt-long recognition element, forming duplex regions that span approximately one third to the full recognition-site length. Because ssDNA and dsDNA differ strongly in their mechanical properties, the recognition element effectively transitions from a flexible ssDNA-dominated to a progressively stiffer polymer chain as the duplex fraction increases. We therefore describe the mixed mechanical behavior of the partially hybridized recognition element by an effective persistence length *L_p_*. Within the worm-like chain (WLC) model, fluctuations in the end-to-end distance are expected to decrease with increasing bending rigidity, *κ* = *k*_B_*TL_p_*. Thus, an increase in effective *L_p_* is reflected in a gradual stiffening of the harmonic potential, leading to reduced angular fluctuations of the lever arm.

While duplex formation determines the overall response, both the angular displacement and fluctuations are also sensitive to the geometry of the entire complex, including the remaining unpaired region of the recognition element (Fig. 2f). Based on equilibrium conformations predicted by NUPACK, the 9, 11, and 13 nt targets adopt distinct loose-end geometries depending on the 5’*→*3’ or 3’*→*5’ binding orientation. For the 11 and 13 nt targets, conformations predicted to contain loose-end hairpins exhibited significantly reduced median Δ*ϕ* values compared with the corresponding hairpin-free conformations (11 nt: 10.9*^◦^* vs. 15.1*^◦^*, *p* = 1.8 *×* 10*^−^*^14^; 13 nt: 12.9*^◦^* vs. 16.0*^◦^*, *p* = 1.7 *×* 10*^−^*^7^, Holm-adjusted; n=55). We attribute this reduction to the mechanical decoupling introduced by the hairpin, which acts as a compliant junction and decreases transmission of the binding-induced contraction to the lever arm. Consistent with this interpretation, hairpin-containing geometries also exhibited smaller Δ*σ* values (Supplementary Fig. S8), indicating reduced stiffening upon target binding. The remaining target lengths further support this interpretation. For the 9 nt target, where both binding orientations are predicted to form similar hairpins, median Δ*ϕ* values differed only marginally (8.0*^◦^* vs. 6.5*^◦^*, *p* = 5.1 *×* 10*^−^*^2^). Likewise, for the 15 nt target, where neither orientation forms a hairpin, the responses were indistinguishable (18.3*^◦^* vs. 19.6*^◦^*, *p* = 2.6 *×* 10*^−^*^1^). Together, these results demonstrate that the mechanical readout is governed primarily by local secondary structure rather than binding orientation, enabling discrimination of subtle conformational differences in ssDNA. By programming the recognition-element geometry, the mechanical transduction of the platform can therefore be tailored to specific sequence lengths and secondary-structure motifs.

### Transient binding and strand-exchange dynamics

We next evaluated the platform’s ability to dynamically monitor molecular binding and unbinding events in real time using hybridization of short oligonucleotides as a model system. Hybridization is generally thought to proceed through nucleation of a few contiguous base pairs, followed by rapid zippering of the remaining helix [26–29]. Once a stable nucleation intermediate has formed, the remaining base pairs zip up within microseconds [30]. Consequently, binding of a short 9 nt oligonucleotide is observable as a sharp transition in the angular time trace (Fig.3a).

At room temperature, the resulting 9 bp duplex is only marginally stable. Continuous monitoring of the nanomechanical amplifier over 300 s in the presence of excess target strands (10 nM) therefore revealed stochastic association and dissociation events as discrete transitions in the angular trajectory. Pooling single-molecule events from 95 structures (Fig. 3b) yielded a slightly skewed distribution of binding-induced lever-arm displacements, Δ*ϕ*, whose dominant population was well described by a Gaussian fit with a peak at *µ*_Δ*ϕ*_ = 8.9*^◦^*. This is in good agreement with the peak of the fitted Δ*ϕ* distribution measured for the steady bound-state at high target concentration (200 nM; *µ*_Δ*ϕ*_ = 8.2*^◦^*). The time trace indicates reduced angular fluctuations in the bound state, likely reflecting the increased rigidity of the recognition–target duplex. We therefore analyzed the change in angular variance upon binding (Fig.3c). Based on the peak of the Gaussian-fitted distribution of changes in lever arm fluctuations, binding indeed suppressed lever arm fluctuations (*µ*_Δ*σ*_ = *−*1.3*^◦^*), which closely aligns with the peak value obtained from the steady bound-state measurement (*µ*_Δ*σ*_ = *−*1.0*^◦^*).

**Fig. 3:**
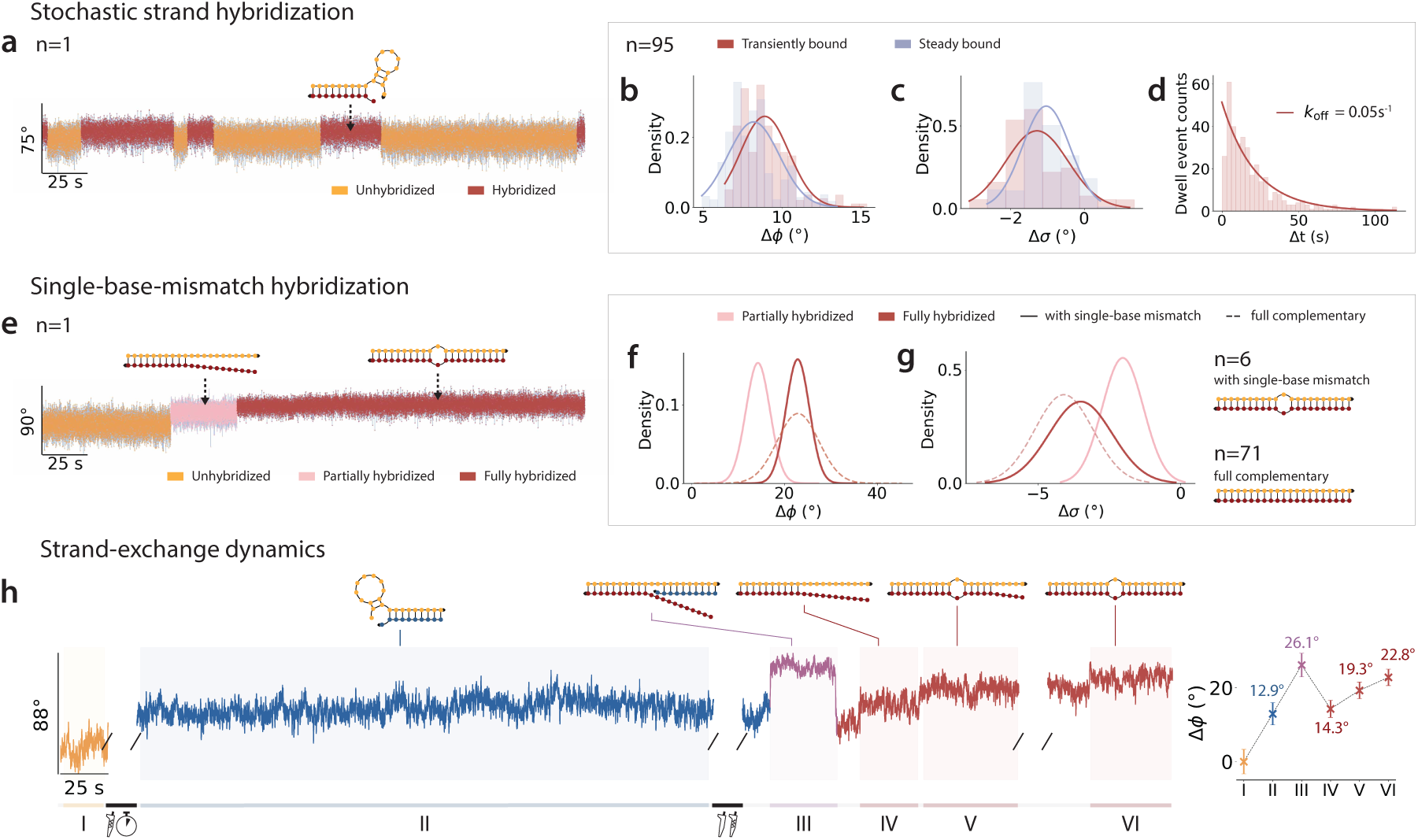
Time-resolved strand hybridization dynamics: transient binding, mismatch recognition and strand exchange. **a**, Single-molecule angular time trace showing stochastic binding and unbinding of a 9 nt target. **b,c,** Overlaid Gaussian curves fitted to Δ*ϕ* and Δ*σ* distributions (bin size = 0.5*^◦^*) comparing the transient and steady bound-state measurements, pooled across 95 structures. **d** Single-exponential fitted dwell time distribution of the transient binding events. **e,** Single-molecule angular time trace showing unhybridized (orange), partially hybridized (pink), and fully hybridized (red) states of a 21 nt target with a central A–A mismatch bound to the recognition strand. **f,g,** Over-laid Gaussian curves fitted to Δ*ϕ* and Δ*σ* distributions comparing single-base mismatch (n=6) and fully complementary target (n=71) hybridization measurements. **h,** Left: Denoised single-molecule trace capturing strand exchange dynamics on a single recognition strand: I, free strand; II, 9 nt bound; III, co-occupancy by 9 nt and 21 nt single-base-mismatch; IV–VI, progressive hybridization of the mismatch target following 9 nt dissociation, yielding a stable bound state (putative strand interactions inferred from Δ*ϕ* analysis are illustrated in the schematics above each trace; detailed protocol in Methods). Right: Segment-specific Δ*ϕ* values referenced to the free strand baseline.

We next analyzed the bound-state dwell-time distribution to quantify the dissociation kinetics. Dwell-time analysis of the bound state reveals single-exponential unbinding kinetics, with *k*_off_ = 0.05 s*^−^*^1^ (Fig. 3d). Our nanomechanical platform thus provides kinetic resolution that is comparable with previous single-molecule FRET [31] and optical trap [32] measurements (0.05–0.1 s*^−^*^1^) under similar buffer conditions (20 mM Mg^2+^).

We further probed the system’s capability to resolve binding dynamics using a 21 nt oligonucleotide containing a single central A–A mismatch. While most trajectories recorded at a 5 ms frame rate exhibited two-state behavior (unbound and bound), a small subset revealed a distinct third state, similar to that shown in Fig. 3e. These three-state trajectories are consistent with the presence of a transient intermediate corresponding to partial duplex formation, which stalls at the mismatch position, but is otherwise obscured by its short lifetime and measurement noise.

Gaussian fits to Δ*ϕ* distributions pooled from trajectories of six structures yielded two distinct populations (Fig. 3f), with one peak matching the fully bound duplex (*µ*_Δ*ϕ*_ = 22.9*^◦^*) and the second peak occurring at *µ*_Δ*ϕ*_ = 14.3*^◦^*, approximately halfway between the unbound and fully bound states, consistent with a partially hybridized intermediate. Lever-arm fluctuations decreased progressively from the unbound to the intermediate and fully bound states (*µ*_Δ*σ*_ = *−*2.0*^◦^* and *−*3.5*^◦^*, respectively; Fig. 3g).

Then, we investigated whether our system could directly monitor the dynamics of DNA strand exchange, a process that is widely used in dynamic DNA nanotechnology [33], and that also plays important roles in numerous biological processes [34, 35]. To this end, we first introduced a 9 nt target strand complementary to the 3*^′^*-terminal of the 21 nt recognition strand. This resulted in a partially occupied recognition strand comprising a 12 nt single-stranded region followed by a 9 bp duplex segment (Fig. 3h, I–II). We then added a 21 nt target containing a single central mismatch. Denoised single-molecule traces (Fig. 3h, III–VI) captured the strand exchange process in real time: following invasion of the exposed receptor ‘toehold’ region by the 21 nt target, a transient state was observed in which both targets were simultaneously bound. This intermediate was followed by dissociation of the 9 nt strand and progressive hybridization of the longer target along the receptor, ultimately yielding a stably bound duplex state. Segment-specific analysis reveals distinct angular displacements (Δ*ϕ*) for partially occupied and stably bound configurations relative to the free-strand baseline, providing a direct, calibration-free single-molecule readout of strand-exchange dynamics that complements previous observations obtained with magnetic or optical tweezers [36, 37]. With >100 individual platforms simultaneously observed within a single field of view, our approach enables parallel single-molecule monitoring of strand-exchange dynamics across multiple structures (see Supplementary Fig. S9).

### Single-molecule readout of transient nanometer-scale aptamer switching

Finally, we sought to challenge our platform with a structural transition that extends beyond simple DNA duplex formation and occurs at the one-nanometer length scale. This regime is central to structure-switching aptamers, which transduce the binding of low-molecular-weight analytes into small conformational rearrangements [38–40]. Such aptamers have previously enabled electrochemical sensors for real-time, *in situ* monitoring of therapeutic agents, metabolites, and neurotransmitters [41, 42]. In such sensors, however, receptor switching is read out through ensemble-averaged electrical signals that report the consequences of conformational change, rather than the structural transition itself [43]. We therefore asked whether our platform could move beyond ensemble electrical readouts and directly resolve the switching dynamics of individual molecules. To address this, we employed a dopamine-binding DNA aptamer reported by Nakatsuka *et al.* [40] that folds into a compact G-quadruplex for target binding, reducing its effective molecular size by only approximately one nanometer (Fig. 4a) [43, 44]. The resulting contraction pulls the lever arm clockwise, producing a negative angular displacement (Δ*ϕ <* 0, Fig. 4b).

**Fig. 4:**
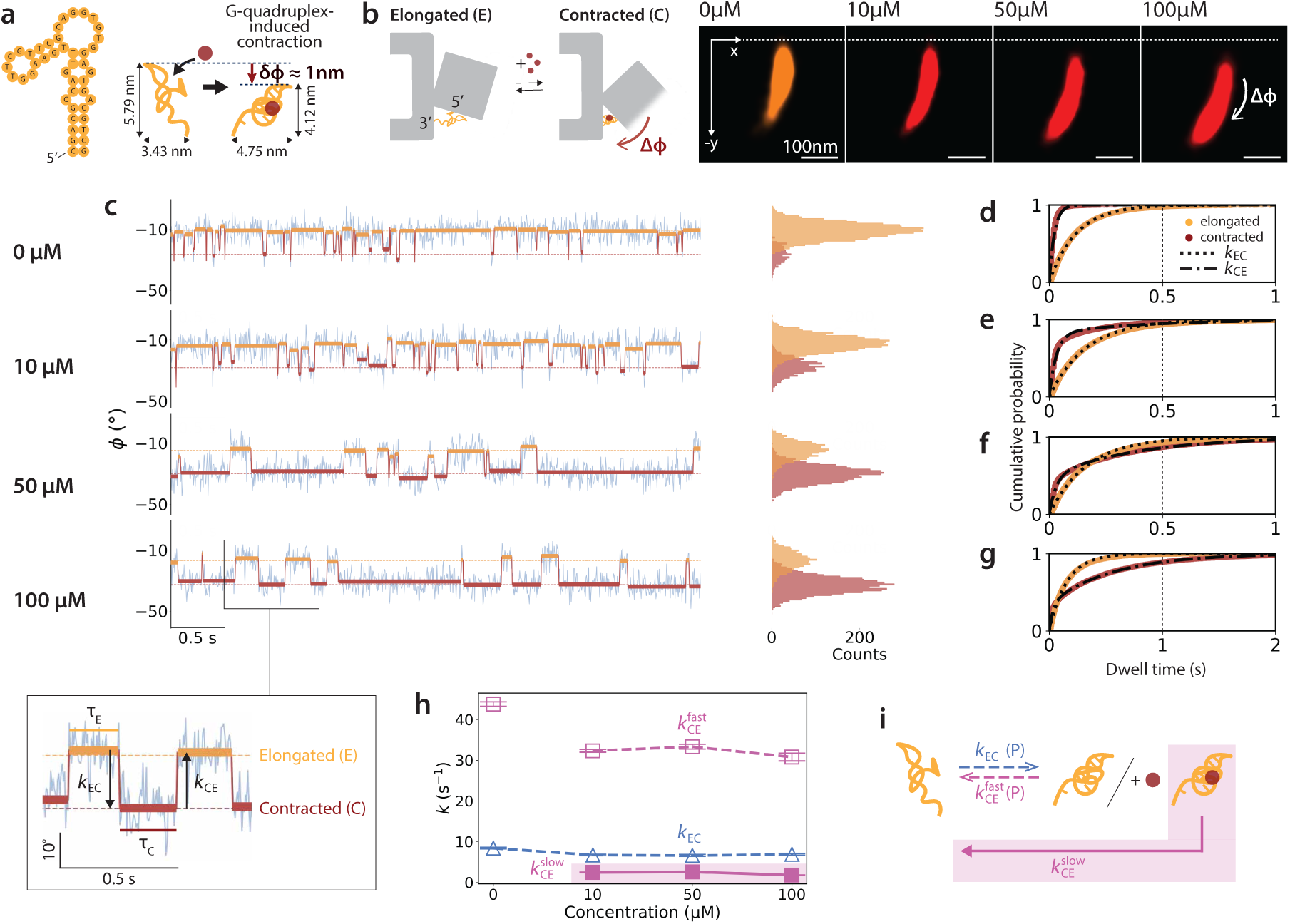
Conformational-selection kinetics of dopamine-aptamer binding. **a**, Schematic illustrating the interconversion of the dopamine-binding aptamer between an elongated and a G-quadruplex-induced contracted conformation, which is selectively stabilized by dopamine binding. **b,** Contraction of the aptamer drives a clockwise displacement of the lever arm (Δ*ϕ <* 0). TIRFM localization heatmaps showing lever arm motion during the first 25 s in the absence of dopamine (orange) and over 120 s at increasing dopamine concentrations (10, 50 and 100 µM, red). **c,** Left: Single-molecule angular time traces of the same platform (blue) showing transitions between the elongated (orange) and contracted (red) states of the aptamer across a dopamine concentration series. Enlarged view highlights individual state dwell times (*τ_E_*, *τ_C_*) and corresponding state transitions (*k*_EC_, elongated *→* contracted; *k*_CE_, contracted *→* elongated). Right: Corresponding HMM-derived angular histograms of the elongated and contracted states. **d-g,** Empirical cumulative distribution function (ECDF) analysis of dwell times for both elongated and contracted states (n=91). **h,** Transition rates across 0, 10, 50, and 100 µM dopamine. Error bars represent 95% confidence intervals (5000 bootstrap iterations). **i,** Schematics summarizing the observed kinetic pathways.

To maximize the angular readout induced by dopamine binding, a truncated version of the aptamer was incorporated into the nanomechanical amplifier [40]. We monitored binding dynamics over a dopamine concentration series spanning 0–100 µM and analyzed the angular time traces using hidden Markov modeling (HMM), revealing two distinct states corresponding to elongated and contracted aptamer conformations with an angular separation of |Δ*ϕ| ≈* 18*^◦^* (Fig. 4c). For the 300 nm lever arm, this angular displacement corresponds to a tip displacement of approximately 94 nm and thus to nearly 100-fold geometric amplification.

The clear separation of the two conformational states allowed us to directly quantify their switching kinetics by dwell-time analysis (Fig. 4d-g, Supplementary Fig. S10), from which we extracted the transition rates in Fig. 4h and established the kinetic scheme shown in Fig. 4i. Interestingly, as exemplified in Fig. 4c, even in the absence of dopamine the aptamer exhibited transient excursions into the contracted state that rapidly returned to baseline, corresponding to a fast contracted-to-elongated transition rate of *k*_CE_ = 43.8 s*^−^*^1^. This suggests that the aptamer stochastically samples a short-lived G-quadruplex conformation even without ligand, whereas dopamine binding shifts the conformational equilibrium toward the contracted state. Consistent with this picture, the contracted-state kinetics showed a fast component *k*_CE,fast_ *≈* 32 s*^−^*^1^, similar to the ligand-free excursions, and a slow component *k*_CE,slow_ *≈* 2.3 s*^−^*^1^ associated with dopamine-induced stabilization of the G-quadruplex (Fig. 4h). In contrast, the closely related precursor L-3,4-dihydroxyphenylalanine (L-DOPA) lacked the slow kinetic component observed for dopamine, indicating that the contracted state is selectively stabilized by the target molecule (Supplementary Fig. S11).

The forward transition from the elongated to the contracted state, meanwhile, remained dominated by a single kinetic component and showed little dependence on dopamine concentration, with *k*_EC_ *≈* 6.7 s*^−^*^1^. Together with the similar angular displacements Δ*ϕ* of the short- and long-lived contracted states (Supplementary Fig. S12), this indicates that dopamine binding primarily modulates the lifetime of the contracted conformation rather than creating a new structural state or substantially altering the G-quadruplex folding rate. These observations are consistent with a conformational-selection mechanism in which the aptamer interconverts between elongated and contracted conformations (*E* ⇌ *C*), while dopamine selectively binds and stabilizes the contracted state (*C* + *D* ⇌ *C*:*D*) [45–49].

Beyond this population-level picture, the single-molecule readout of our platform provided access to molecular heterogeneity that is typically obscured in ensemble-averaged measurements. Individual aptamers separated into kinetically distinct subpopulations (Supplementary Fig. S13), and these differences persisted upon dopamine binding. This highlights substantial kinetic heterogeneity within the aptamer ensemble and suggests that individual molecules sample distinct regions of a rugged conformational energy landscape, even though dopamine stabilizes the contracted state in a broadly similar manner across the population.

Finally, we also tested a second, serotonin-binding aptamer that is reported to undergo a G-quadruplex-associated conformational change analogous to the dopamine-binding aptamer [40]. Although its overall architecture is similar, it exhibited larger angular fluctuations and no clearly discernible conformational states in the reference measurements (Supplementary Fig. S14). We speculate that this reflects a less stable stem that is more susceptible to fraying than that of the dopamine-binding aptamer. These differences suggest that the mechanical readout of our platform depends not only on the magnitude of the conformational transition, but also on the intrinsic stability of the aptamer architecture.

Our results demonstrate that the platform can monitor binding-coupled conformational dynamics beyond nucleic acid hybridization at the level of individual aptamer molecules, despite the inherently small length scales and rapid fluctuations associated with such transitions. At the same time, the comparison between aptamers shows that the accessible dynamic range depends on the structural stability of the receptor, particularly the aptamer stem. By resolving the behavior of individual receptors rather than ensemble-averaged responses, the platform provides access to molecular heterogeneity, switching kinetics, and structural constraints that are otherwise difficult to disentangle. More broadly, this single-molecule resolution of our platform uncovers properties relevant to aptamer selection and design that remain hidden in conventional ensemble-averaged readouts.

## Conclusion

By leveraging the programmability of DNA origami, we establish a nanomechanical sensing platform that transduces molecular interactions into amplified mechanical signals, enabling direct quantification of nanoscale conformational changes with single-molecule sensitivity without requiring analyte labeling. The structural precision and modularity of the platform facilitate the integration of diverse recognition elements. In principle, the same framework could be extended to monitor protein–nucleic acid interactions, nucleic acid processing by enzymes, or other biomolecular reactions that generate a mechanical or conformational response. As demonstrated here for DNA hybridization, DNA conformational transitions, DNA strand displacement reactions, and aptamer–target binding, the approach supports time-resolved monitoring of a broad range of molecular interactions. Beyond these examples, the same framework could be extended to protein–nucleic acid interactions, enzymatic processing of nucleic acids, and other biomolecular reactions that couple molecular activity to a conformational response.

The ability to resolve subtle geometric variations, transient binding events, and heterogeneous kinetic pathways highlights its potential for interrogating complex molecular processes beyond the capabilities of conventional fluorescence-based measurements. Mechanical signal transduction circumvents reliance on intensity-based readouts and calibration-sensitive distance measurements. Furthermore, the fluorescence labels used here for motion tracking could readily be replaced by nanoparticle-based probes, enabling continuous observation without photobleaching and reducing susceptibility to photophysical artifacts.

More broadly, we anticipate that mechanically amplified angular readouts could provide a scalable framework for receptor-target fingerprinting. Coupled with AI-assisted analysis, such signals may enable the identification and classification of molecular targets based on their characteristic dynamic and conformational signatures. Integration with controlled surface-patterning strategies, such as the lithographic arrangement of DNA origami platforms with distinct receptors in defined orientations [50], could further enable the generation of collective signal patterns that serve as multiplexed barcodes for the identification of complex analyte mixtures. The current random immobilization strategy requires low surface densities to avoid overlap between neighboring platforms, which in turn necessitates elevated ligand concentrations to observe sufficient binding events within practical measurement times. Controlled positioning of sensing platforms could substantially increase usable surface densities and thereby improve sensitivity, potentially enabling operation at detection limits approaching those of conventional biosensors. Combined with emerging stabilization strategies for DNA origami, including covalent crosslinking [51] and silicification [52], the platform may ultimately become compatible with complex biological environments such as blood plasma. More generally, these developments point toward nanomechanical sensing architectures that connect molecular recognition and conformational dynamics to amplified, information-rich mechanical signals for future biosensing and diagnostic applications.

## Methods

### Design of DNA origami structures

DNA origami structures were designed using cadnano 2.4.13 [53], with validation of three-dimensional geometrical features and structural integrity performed using oxDNA simulations [54].

### Folding of DNA origami structures

DNA origami structures were folded using a one-pot self-assembly strategy by mixing 30 µl of 100 nM p8064 scaffold (produced in-house and stored in ddH_2_O at -22*^◦^*) with 5-fold molar excess of basic origami structure staples and 10-fold molar excess of system- and target-specific modified staples. LNA- and ATTO fluorescent dye-modified strands were purchased from biomers.net GmbH in 0.1xTE buffer, pH 8.0, whereas all other strands were purchased from Integrated DNA Technologies (IDT) in 100 µM IDTE buffer, pH 8.0. Complete staple sequences are listed in Supplementary Table S1-S4. The supporting base, forming a compliant hinge with the lever arm (hereafter referred to as the base monomer), and the lever arm extension (hereafter referred to as the extension monomer) were folded separately in individual self-assembly reactions.

Folding reactions were carried out in a folding buffer (containing 50 mM Tris-Base, 10 mM EDTA, 50mM NaCl, and 20 mM MgCl_2_). Thermal annealing was performed using a peqSTAR thermocycler (VWR Peqlab) with a gradual temperature ramp from 70*^◦^*C to 40*^◦^*C, decreasing at a rate of 0.1*^◦^*C every 2 minutes. Upon completion of the annealing program, folded structures were stored at 20*^◦^*C until further use.

### Post-folding DNA origami purification and dimer assembly

Folded DNA origami structures were purified to remove excess staple strands using polyethylene glycol (PEG) precipitation. An equal volume of PEG buffer matching the folding solution (containing 15%PEG, 5 mM Tris-Base, 1 mM EDTA, 50.5 mM NaCl, and 20 mM MgCl_2_), was added to the reaction mixture. The mixture was centrifuged at 20*^◦^*C and 20,000 rcf for 20 minutes using an Eppendorf Centrifuge 5424 R. Faint white pellets of the folded structures formed on the tube walls, and the supernatant was carefully removed. Pellets were resuspended in 20 µl of storing buffer (containing 5 mM Tris-Base, 1 mM EDTA, 5 mM NaCl, and 20 mM MgCl_2_). The concentration of the resuspension mixture was determined using an Implen NanoPhotometer N6.

For base monomers, a 100-fold molar excess of NeutrAvidin per biotinylated staple relative to the measured concentration was added and incubated for 10 minutes at room temperature. For extension monomers, a 10-fold molar excess of ATTO655-modified LNA strands per handle-sequence-modified strand was added and incubated for 1 hour at 37*^◦^*C in an Eppendorf ThermoMixer Comfort.

The mixed solutions were further purified by agarose gel electrophoresis. A 2% (w/w) agarose gel was prepared in running buffer (containing 0.5*×*TBE and 5.5 mM MgCl_2_). Samples were mixed with 16.6% (v/v) viscous loading dye (containing 0.05% bromophenol blue, 40% sucrose, 0.1 M EDTA pH 8.0, and 0.5% SDS) and loaded into the gel wells separately. Electrophoresis was run for 1 hour at 70 V under constant fan cooling. Folded structures labeled with ATTO488 were visualized under blue light, leading bands that were sharp and bright, indicative of well-folded origami structures, were excised from the gel. Final concentration of the purified structures was measured using the NanoPhotometer.

Purified base and extension monomers were incubated at a 1:1 stoichiometric ratio in an Eppendorf ThermoMixer Comfort at 37*^◦^*C for 1 hour to assemble the final dimer platform.

### Sample preparation for TIRFM measurements

All TIRFM measurements were performed using sticky-Slide VI 0.4 flow chambers (ibidi) assembled with biotinylated PEG-passivated glass coverslips. These PEG-biotin-coated coverslips were prepared according to established protocol [21]. Unless otherwise stated, all slide preparation, structure dilution, and measurement steps were carried out in imaging buffer (containing 5 mM Tris-Base, 1 mM EDTA, 5 mM NaCl, and 20 mM MgCl_2_), supplemented with 0.05% (v/v) Tween to minimize nonspecific surface interactions.

Prior to sample introduction, coverslips were hydrated with 200 µl ddH_2_O and rinsed once with 200 µl imaging buffer. Samples were diluted to 50 pM in imaging buffer, pipetted into the flow chambers, and incubated for 2 minutes to enable surface immobilization. The flow chamber was then washed three times with 200 µl imaging buffer. After washing, 40 µl imaging buffer was pipetted into the flow chamber to prevent drying-induced structural unfolding.

### TIRFM acquisition and measurement protocol

All TIRFM measurements were performed using MMStudio (Micro-Manager v2.0.0-gamma1). Image acquisition was carried out with an exposure time of 5 ms, and the total number of frames was adjusted according to the duration of each measurement.

For strand hybridization measurements shown in Fig. 2, all measurements were performed at a target strand concentration of 200 nM to achieve effective saturation for shorter strands that otherwise bind transiently. Before target introduction, a 25 s reference measurement was recorded to establish the baseline fluctuations of the lever arm in the absence of target. This reference quantified intrinsic mechanical noise and confirmed stable surface immobilization. The imaging buffer was then exchanged with buffer containing 200 nM target strand and allowed to equilibrate for 5 minutes. Subsequent target-bound measurements were recorded for 25 s. For each target length, measurements were performed in separate flow chambers using freshly prepared structures.

All measurements shown in Fig. 3 have first a 25 s reference measurement recorded followed by distinct measurement protocols after target introduction. For 9 nt target measurement, the imaging buffer was exchanged with buffer containing 10 nM target strand and allowed to equilibrate for 5 minutes, followed by a 5-minute measurement. For single-base mismatch target measurement, the imaging buffer was exchanged with buffer containing 10 nM target strand and a 5-minute measurement was initiated immediately. For strand-exchange measurements, the imaging buffer was first exchanged with buffer containing 200 nM 9 nt target strand and allowed to equilibrate for 5 minutes, followed by a 5-minute measurement. The flow chamber was then washed three times with 200 µl imaging buffer, and filled with buffer containing 10 nM single-base mismatch target strand. Subsequent measurements were started immediately and continued until photo-bleaching limited further imaging.

All measurements shown in Fig. 4 were conducted exclusively in an aptamer-optimized imaging buffer (containing 5 mM Tris base, 1 mM EDTA, 5 mM NaCl, 5 mM MgCl_2_, 100 mM KCl and 1 mM CaCl_2_, supplemented with 0.05% (v/v) Tween) stored at 4*^◦^*C prior to use. A 25 s reference measurement was recorded prior to target introduction. Dopamine target solutions were prepared just before the target measurements owing to the rapid oxidative polymerization of dopamine in aqueous solution. Dopamine powder was dissolved directly in imaging buffer to obtain a 100 µM stock solution, with the buffer volume adjusted according to the weighed mass. Solutions of 50 µM and 10 µM dopamine were prepared by further dilution of this stock solution in separate tubes. Target measurements were performed sequentially at increasing target concentrations within the same flow chamber. For each concentration step, buffer exchange was followed by 5-minute equilibration and acquisition for 120 s. Between each concentration increment, three washing steps with 200 µl imaging buffer were performed to remove residual ligand.

For serotonin measurements, a 100 µM serotonin target solution was prepared similarly. A 25 s reference measurement was recorded prior to target introduction, followed by acquisition of the 100 µM serotonin target measurement for 120 s after buffer exchange and a 5-minute equilibration. Dopamine and serotonin powders were purchased from Merck/Sigma-Aldrich.

### Image processing and data analysis

Localization of single-molecule fluorescence signals and generation of TIRFM localization heatmaps for selecting particles of interest were performed using the Picasso v0.7.3 software package [55]. Data from sequential measurements were overlaid and drift-corrected. Localization clusters corresponding to the same structure were manually picked and annotated with particle number. Only localization clusters exhibiting distinct circular trajectories spanning *≤* 90*^◦^*, consistent with the geometric constraints of the structural design and free from overlapping localizations originating from neighboring structures, were picked. All picked localizations were exported for Python-based angular analysis.

Conversion of particle localizations to polar coordinates was performed using custom Python-based analysis scripts [56]. For each picked particle, circle fitting was done on all underlying localizations collectively. Angular coordinates (*ϕ*) were calculated from the x- and y-coordinates relative to the fitted circle center and wrapped to the interval (*−π, π*] to account for boundary effects. To enable cross-particle analysis, each particle was reoriented by aligning the 99th percentile of the angular distribution from the 25 s reference to 0°, with all other angular positions re-referenced accordingly. Particles with a fitted radius outside the range of 300 *±* 50 nm were excluded from further analysis due to issues with circle fitting.

For the results shown in Fig. 2, mean lever arm positions in target-free 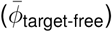 and target-bound 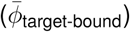 trajectories were calculated from the circular means of the corresponding angular distributions to address the angular boundary problem. Difference between the circular mean values computes the target-binding induced lever arm displacement, given by 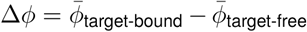.

For the results shown in Fig. 3 and Fig. 4, HMM analysis was applied to the angular trajectories to identify distinct segments associated with target-free and target-bound angular states. For stochastic strand hybridization (Fig. 3a–d) and target–aptamer binding measurements (Fig. 4), the angular trajectories were analyzed using a two-state Gaussian HMM, whereas a three-state Gaussian HMM was used for the single-base-mismatch strand hybridization measurements (Fig. 3e–g). Initial state means and variances were obtained using K-means clustering with two or three clusters, as appropriate, using a fixed random seed of 42. Initial state probabilities and transition probabilities were set uniformly. A Gaussian HMM with diagonal covariance matrices was subsequently fitted to each angular trajectory for up to 200 iterations, with a convergence tolerance of 10*^−^*^3^. The most probable hidden-state sequence was then obtained from the fitted model. As HMM state labels were assigned arbitrarily, all fitted states were reordered according to the mean angle calculated for each state, such that state 0 corresponded to the lowest mean angle, state 1 to the higher mean angle, while for the three-state model, state 2 corresponded to the highest mean angle. All HMM fitting parameters, including the fitted emission means and variances, initial state probabilities, and transition matrices, were stored for each individual platform and retained for subsequent state prediction where required. For the strand hybridization measurements shown in Fig. 3a-g, an additional state-sequence refinement was applied to merge short contiguous segments with neighboring segments when the mean angle of the short segment was close to that of the neighboring segments. This post-processing step was applied to mitigate apparent state switching caused by noise or HMM misclassification while retaining sustained transitions between angular states. Subsequently, the mean and standard deviation of the angular trajectory were calculated for each state. All state-specific data were stored in a second dataframe for further analysis.

Single-molecule Δ*ϕ* responses were pooled across all platforms acquired under the same measurement condition. The resulting distributions were visualized as swarm plots with overlaid box plots indicating the median (center line), interquartile range (IQR; box), and whiskers extending to xIQR. The slope per two-base increment was obtained from a linear regression of the median values across all target lengths. Δ*ϕ* distributions shown in Fig. 3 were fitted with a Gaussian model to extract the peak values (*µ*_Δ*ϕ*_) for each dataset.

Lever arm fluctuations were quantified by calculating the standard deviation of the target-free (*σ*_target-free_) and target-bound (*σ*_target-bound_) trajectories. In the transient binding regime, *σ* was determined from the corresponding trajectory segments with the longest lifetime. Binding-induced stiffening of the lever arm was quantified as the change in fluctuation amplitude between the target-bound and target-free states, defined as Δ*σ* = *σ*_target-bound_ *− σ*_target-free_.

Single-molecule Δ*σ* responses were pooled across all platforms acquired under the same measurement condition. For each target length shown in Fig. 2e, Supplementary Fig. S7, and Supplementary Fig. S8, the distribution of Δ*σ* values was summarized by the median values, with error bars representing the half IQR of the corresponding distributions, calculated as (*error* = (*Q*_3_ *− Q*_1_)*/*2). For Fig. 3, Δ*σ* distributions were fitted with a Gaussian model to extract the peak values (*µ*_Δ*σ*_) for each dataset.

### Statistical analysis

For every target strand length, Δ*ϕ* values from the two binding orientations were compared using two-sided Welch’s unpaired t-tests. Effect sizes were quantified using Cohen’s d. All resulting p-values were adjusted for multiple comparisons using the Holm–Bonferroni procedure implemented in Python (statsmodels) with *α* = 0.05. Adjusted *P <* 0.05 was considered statistically significant.

### Dwell time analysis

Dwell time distribution of transient 9 nt binding events was first filtered to remove “censored” events arising from the start and end of the measurement window. The remaining dwell-time histogram was subsequently fitted with a single-exponential decay model, 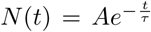, where τ represents the characteristic lifetime of the bound state. The corresponding dissociation rate was calculated from the fitted lifetime using 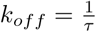 .

Dwell time distributions of dopamine aptamer transitions between the elongated and contracted states (EC: elongated*→*contracted; CE: contracted*→*elongated) were analyzed using cumulative distribution functions (CDFs). Single-exponential 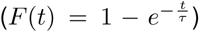 and double-exponential 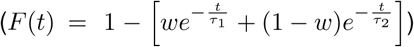 models were evaluated using BIC, with the double-exponential model providing better description of the dwell-time distributions for both transitions; however, the improvement over the single-exponential model was only marginal for the EC transition, in contrast to the substantially better improvement observed for the CE transition. The fitted characteristic lifetimes, *τ*_1_ and *τ*_2_, were converted into kinetic rates using 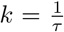 yielding fast (*k*_fast_) and slow (*k*_slow_) transition rates corresponding to the short- and long-lived kinetic components, respectively. Rate uncertainties were quantified by bootstrap resampling of the dwell-time datasets (5000 iterations), followed by CDF refitting for each resampled dataset. The 95% confidence intervals were calculated from the 2.5th and 97.5th percentiles of the resulting bootstrap rate distributions.

### Software and computational analysis

Single-molecule data analysis was implemented using code developed with the assistance of large language models (LLMs), including ChatGPT. Data visualization and manuscript language editing were also assisted by LLMs. All manuscript content was reviewed and verified by the authors.

## Data Availability

Experimental data underlying all figures are available at [repository]. Source data are provided with this paper.

## Code Availability

Custom Python scripts used for single-molecule data analysis are available at [repository].

## Acknowledgments

Funded by the Deutsche Forschungsgemeinschaft (DFG, German Research Foundation) under Germany’s Excellence Strategy – EXC 3092 – 533751719 (Cluster of Excellence Biosystems Design Munich, BioSysteM). This work was additionally funded by the Deutsche Forschungsgemeinschaft (DFG, German Research Foundation) – 201269156 (SFB 1032, project A02).

## Author Contributions

R.Y.L. and E.K. planned the research. E.K., L.J.K.W., and F.C.S. supervised the research. R.Y.L., E.K., and L.J.K.W. developed the DNA nanostructure design rationale. R.Y.L. designed the DNA nanostructures, with E.K. reviewing the design details. R.Y.L. validated the structural integrity using oxDNA simulations. R.Y.L. folded and purified the structures. R.Y.L. performed the single-molecule measurements. R.Y.L. optimized the nanostructure design and fluorescence signal output with guidance from E.K. and L.J.K.W. R.Y.L. performed transmission electron microscopy (TEM) measurements. R.Y.L. developed the single-molecule data analysis pipeline, incorporating a previously developed circle-fitting package optimized by L.J.K.W. E.K. and L.J.K.W provided guidance on data interpretation and advised R.Y.L. on experimental planning. R.Y.L. generated the figures. R.Y.L., L.J.K.W., and F.C.S. wrote the manuscript. All authors discussed and reviewed the manuscript. F.C.S. secured the funding for the project.

## Additional information

**Supplementary information** The online version contains supplementary material.

**Correspondence and requests for materials** should be addressed to Friedrich C. Simmel or Lennart J.K. Weiß.

