## Supplementary Material for "DNA Origami Nanomechanical Amplifiers for Resolving Single-Molecule Binding Events"

**Supplementary references** 36

### Origami design

The nanomechanical amplifier leveraged a lever arm mechanism to convert sub-nanometre conformational changes induced by target binding into a directly measurable fluorescent output. To maximize mechanical amplification by extending the effective lever arm while maintaining structural integrity within a dimeric architecture, we applied a hollow cuboid approach with extensive crossover connections between adjacent helices to mechanically reinforce the structure and minimize local deformation (Supplementary Fig. S1).

In addition to examining the structural integrity of the assembled nanomechanical amplifiers by transmission electron microscopy (TEM), we also employed DNA-PAINT [1] to assess structural integrity under experimental conditions (Supplementary Fig. S2). Specifically, we measured the distance between the hinge region, approximated by binding to a recognition strand, and the distal tip of the lever arm. The experimentally measured distance closely matched the value predicted from the design, confirming that the intended lever-arm geometry was preserved upon assembly.

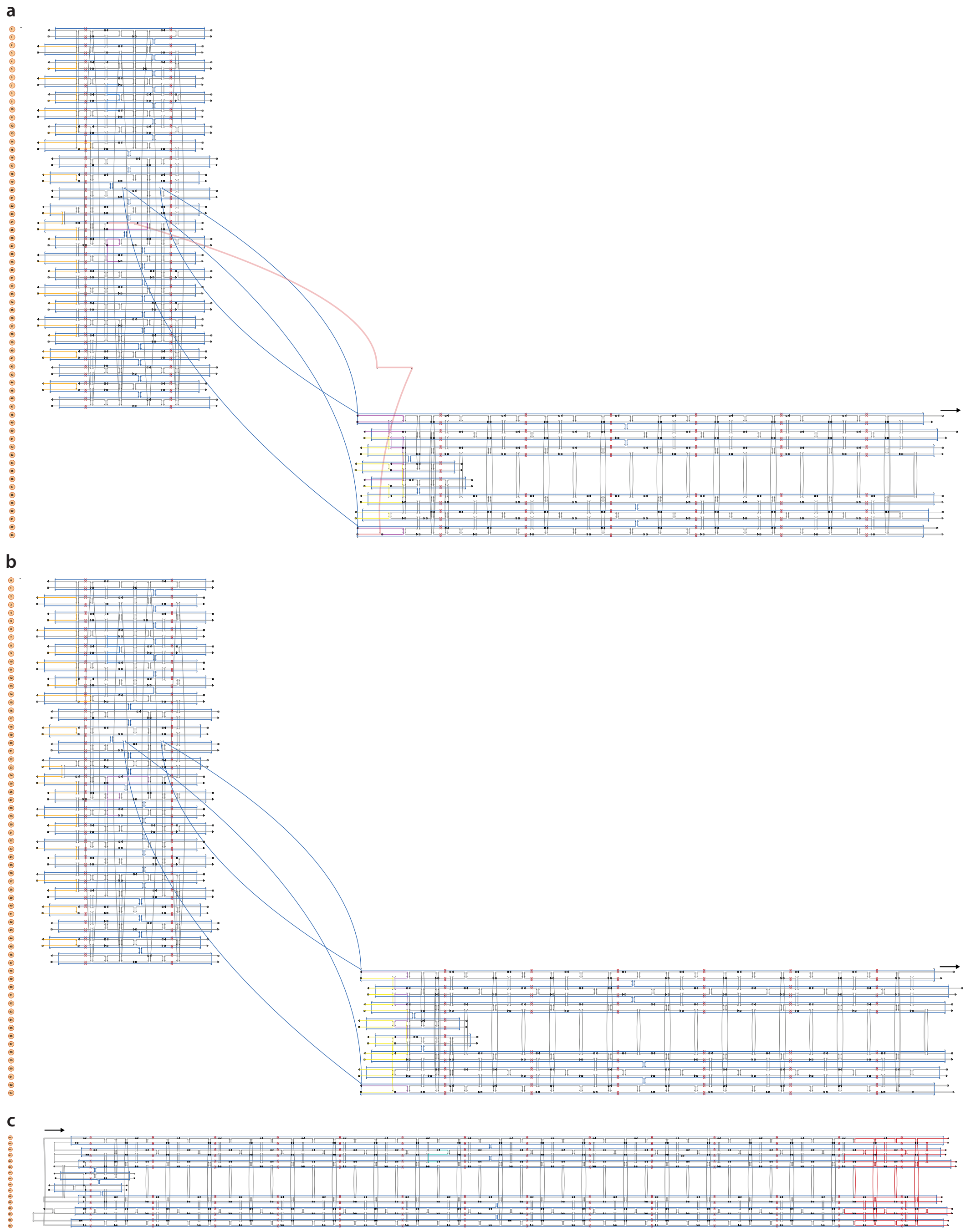

**Fig. S1**: **Cadnano design of the nanomechanical amplifier. a-c,** Cadnano schematics of the two platform base monomer configurations used in the elastic spring (**a**) and steric hindrance (**b**) system, and the lever arm extension monomer (**c**). The arrow indicates the direction of dimeric assembly via shape complementarity to the lever arm extension. The base monomer comprises a supporting base connected to a lever arm through four double-stranded DNA (dsDNA) crossovers that collectively form a compliant hinge. Grey staples indicate unmodified strands used to fold the DNA origami structure. Orange staples denote biotinylated strands modified at either the 5′- or the 3′-end. Cyan staples indicate ATTO488-modified strands used for band excision after gel electrophoresis purification. Peach staple denotes recognition strand tethered between the supporting base and the lever arm to generate spring-like mechanical compliance upon target binding, whereas yellow staples denote recognition strands projecting from the hinge-proximal terminus of the lever arm in the steric hindrance system for target binding. Purple staples indicate the staple variants used to fold each system. Red staples contain handle sequences for hybridization with ATTO655-labeled complementary strands for total internal reflection fluorescence microscopy (TIRFM) measurements.

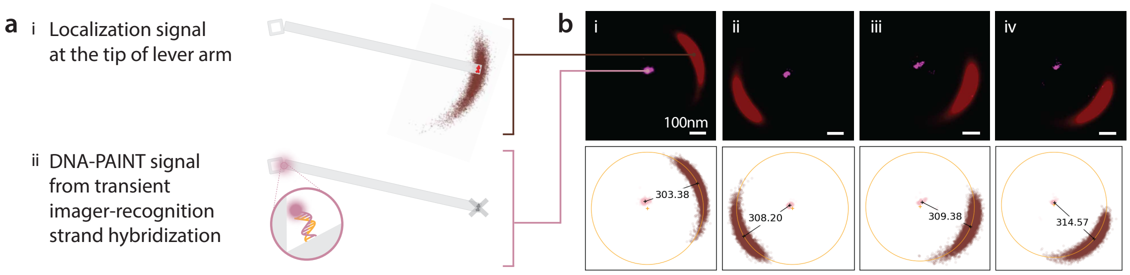

**Fig. S2**: **DNA-PAINT quantification of the lever arm tip-to-hinge distance. a,** Top-view schematic of sterically constrained system, featuring a single DNA-PAINT handle positioned at the hinge-proximal site (see Supplementary Fig.S6**a(ii)**). **(i)** Lever arm trajectories were obtained from permanently attached fluorescent dyes at the lever arm tip, whereas **(ii)** DNA-PAINT signals arose from transient hybridization of ATTO655-labeled imager strands to the recognition-strand handle. Prior to DNA-PAINT acquisition, the lever arm fluorophores were photobleached under high laser power ≈150mW for 30 minutes to prevent signal overlap. All measurements were conducted in imaging buffer containing 45% (v/v) sucrose. **b, i-iv**: Overlaid TIRFM localization heatmaps showing representative single lever arm trajectories (brown) and DNA-PAINT signals (magenta) from four different platforms (top). Localization clusters were fitted to circles to extract angular positions for both the lever arm trajectories and DNA-PAINT signals (bottom). Circular means were calculated for each angular cluster, and the Euclidean distance between the mean positions was used to quantify the separation between the recognition strand and the lever arm tip. Measured distances across four individual platforms were **(i)** 303.4 nm, **(ii)** 308.2 nm, **(iii)** 309.4 nm, and **(iv)** 314.6 nm, consistent with the theoretical distances estimated from the design geometry (307-328 nm).

### Exemplary field of view

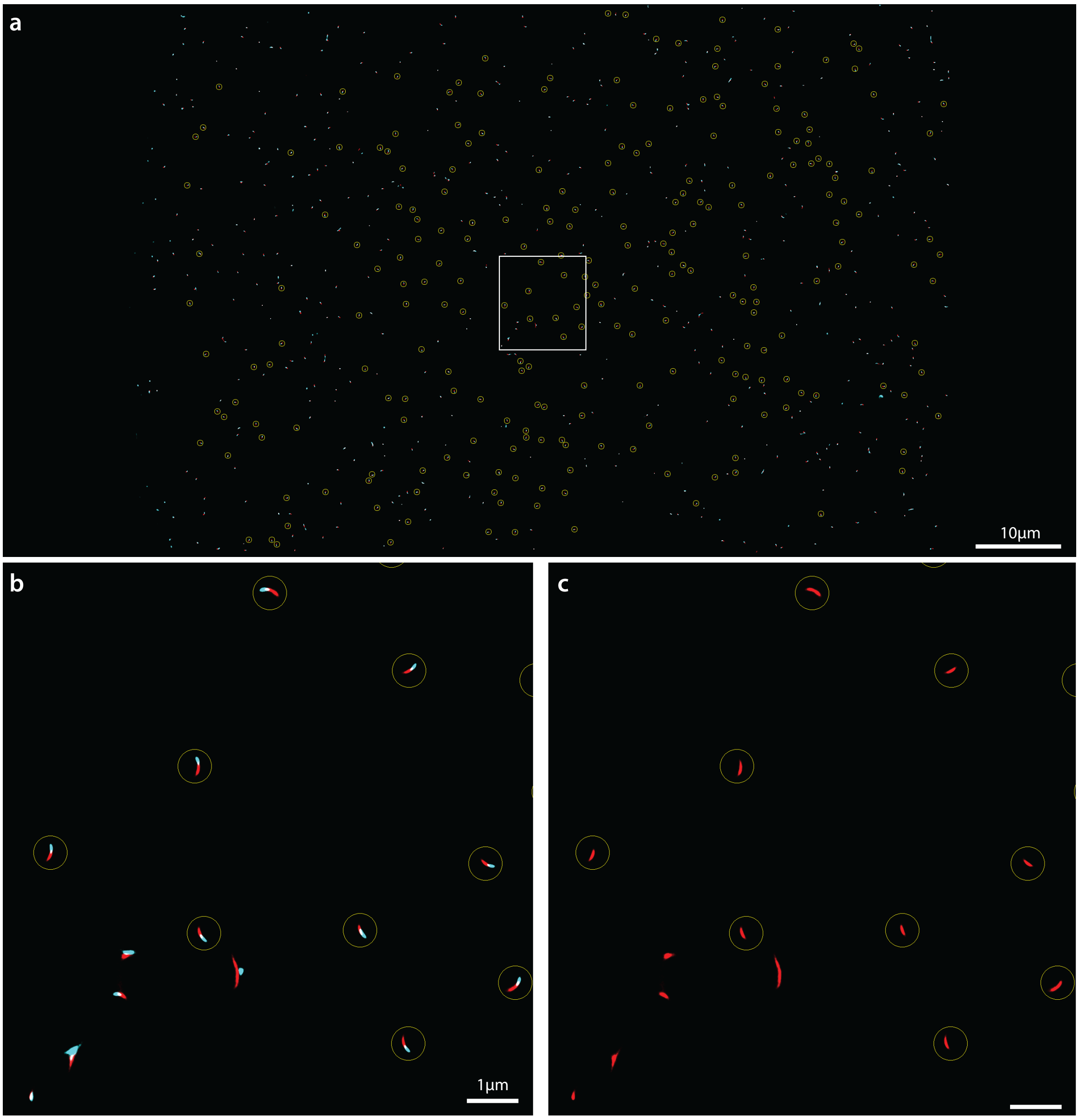

**Fig. S3**: **Exemplary full localization heatmap for 21 nt target strand measurement shown in Fig. 1f. a,** Raw localization data overlaying 25 s target-free measurement (red) and 120 s target-bound measurement (cyan), acquired with a frame time of 5 ms over a field of view of approximately $95 \times 65\ \mu\text{m}^2$. Localization clusters corresponding to the same structure selected for angular analysis are indicated by yellow circles. **b,** Enlarged view of the region enclosed by the white bracket in **a**. Localization clusters were selected based on the target-free measurement, which exhibited distinct circular trajectories spanning $\leq 90°$, while excluding overlapping localizations arising from neighboring structures. **c,** Exemplary target-free localizations corresponding to the localization clusters shown in **b**.

### Lever arm dynamics

Accurate quantification of lever arm motion is essential for sensitively detecting changes in angular trajectories induced by target binding. We therefore first characterized lever arm motion under free diffusion and electric-field actuation and compared the observed range of motion with that predicted by the structural design (Supplementary Fig. S4a,b). We further assessed whether the measurement conditions faithfully resolved the instantaneous angular position of the lever arm or instead reported a temporally averaged position by measuring the lever arm free diffusion across different buffer viscosities and acquisition frame rates (Supplementary Fig. S4c). In DNA-origami-based systems, $Mg^{2+}$ concentration can also substantially affect structural integrity and, consequently, lever-arm dynamics. We therefore characterized lever arm free diffusion at varying buffer $Mg^{2+}$ concentrations (Supplementary Fig. S4d).

For electric-field actuation, measurements were performed in 100 mM NaCl buffer containing 48% sucrose at a frame rate of 2 ms. Lever arm motion was first recorded under free diffusion for 10 s, followed by application of electric fields of 50, 80, 100, 120, and 150 V for 5 s each, with the field applied sequentially over $360°$ to actuate the lever arm. The lever arm was then allowed to return to free diffusion for the remaining 5 s of the measurement. Electric-field actuation enabled the lever arm to explore the full geometrically permitted angular range, whereas free diffusion remained confined to a narrower angular window imposed by the hinge architecture. The range of motion (ROM) was quantified using a percentile-based metric, $\mathrm{ROM} = P_{99.9\%} - P_{0\%}$, to minimize the influence of extreme outliers, which presumably arose from overbending of the lever arm around the base during electric-field actuation. Free-diffusing lever arms exhibited a $\mathrm{ROM}_{\mathrm{FD}} \approx 64.7°$, whereas electric-field actuation yielded a $\mathrm{ROM}_{\mathrm{EF}} \approx 81.1°$. The electrically actuated ROM was consistent with the geometrically designed range of lever arm motion, whereas, as expected, free diffusion sampled a reduced angular range owing to constraints imposed by the hinge.

We next examined lever arm motion under different buffer viscosities and measurement frame rates. The corresponding angular distributions are shown for each condition, with mean angular positions, denoted $\bar{\phi}$, of $-14.9°$ (no sucrose, 5 ms frame rate), $-20.5°$ (45% sucrose, 5 ms frame rate), and $-24.7°$ (45% sucrose, 2 ms frame rate). These measurements reveal the effects of fluctuation timescales and temporal averaging on the measured angular position. Acquisition at 5 ms frame rates introduced substantial temporal averaging of the angular readout but enabled sampling of a broader field of view containing more than 100 structures. By contrast, 2 ms acquisition resolved faster fluctuations but limited particle sampling to fewer than 20 structures owing to hardware constraints. Sucrose-containing buffers reduced temporal averaging, but the resulting increase in viscosity may affect target-binding efficiency.

Finally, we examined changes in lever arm motion with increasing $Mg^{2+}$ concentration in the buffer. Angular distributions compiled from 22 platforms across all conditions are shown, with $\bar{\phi}$ values of $-15.9°$ (5 mM $Mg^{2+}$), $-8.9°$ (12 mM $Mg^{2+}$) and $-4.9°$ (20 mM $Mg^{2+}$), relative to the 100 mM $Na^{+}$ reference condition. Increasing $Mg^{2+}$ concentration produced a progressive shift from a more open hinge configuration, characterized by more negative $\bar{\phi}$ values, towards a more closed configuration with less negative $\bar{\phi}$ values.

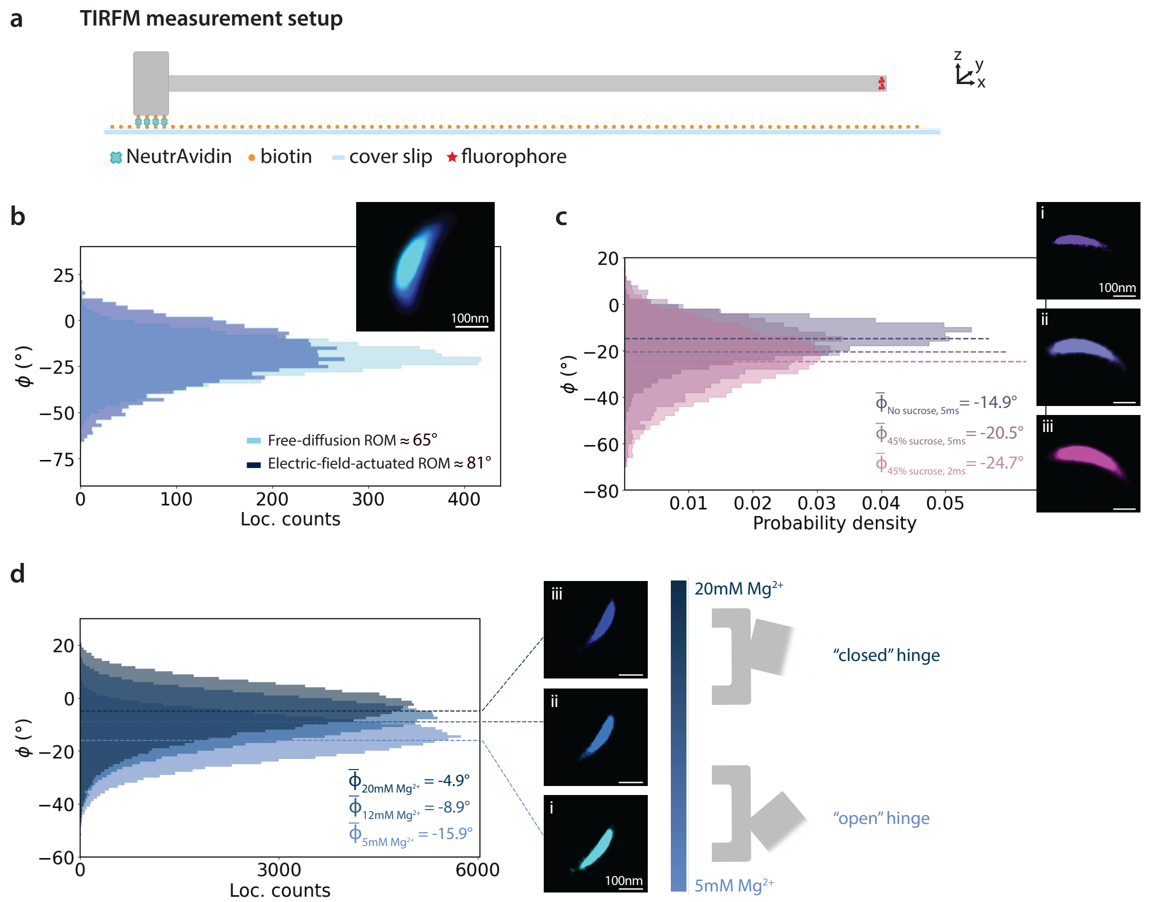

**Fig. S4**: **Structural properties and lever arm dynamics. a,** Side-view schematic of the TIRFM measurement setup. The platform is immobilized on the coverslip via biotin-NeutrAvidin coupling. **b,** TIRFM localization heatmap showing single lever arm motion under $360°$ electric-field actuation (blue), overlaid with trajectories in free diffusion condition (cyan). Corresponding angular distributions of lever arm trajectories under free diffusion and electric-field-controlled conditions are shown. **c,** TIRFM localization heatmaps showing single lever arm motion over 5 s under three measurement conditions: **(i)** 5 ms frame rate in 100 mM $Na^{+}$ buffer without sucrose, **(ii)** 5 ms frame rate in 45% sucrose buffer, and **(iii)** 2 ms frame rate in 45% sucrose buffer. Corresponding angular distributions of lever arm trajectories are shown for each condition. **d,** TIRFM localization heatmaps showing single lever arm motion over 5 s at increasing $Mg^{2+}$ concentrations (**(i)** 5 mM, **(ii)** 12 mM, and **(iii)** 20 mM). Angular distributions of the lever arm trajectories compiled from 22 platforms across all conditions, relative to the 100 mM $Na^{+}$ reference condition (not plotted), are shown. Top-view schematics of the platform illustrate a progressive shift from a more open to a more closed configuration with increasing $Mg^{2+}$ concentration, as indicated by increasing less negative $\bar{\phi}$ values.

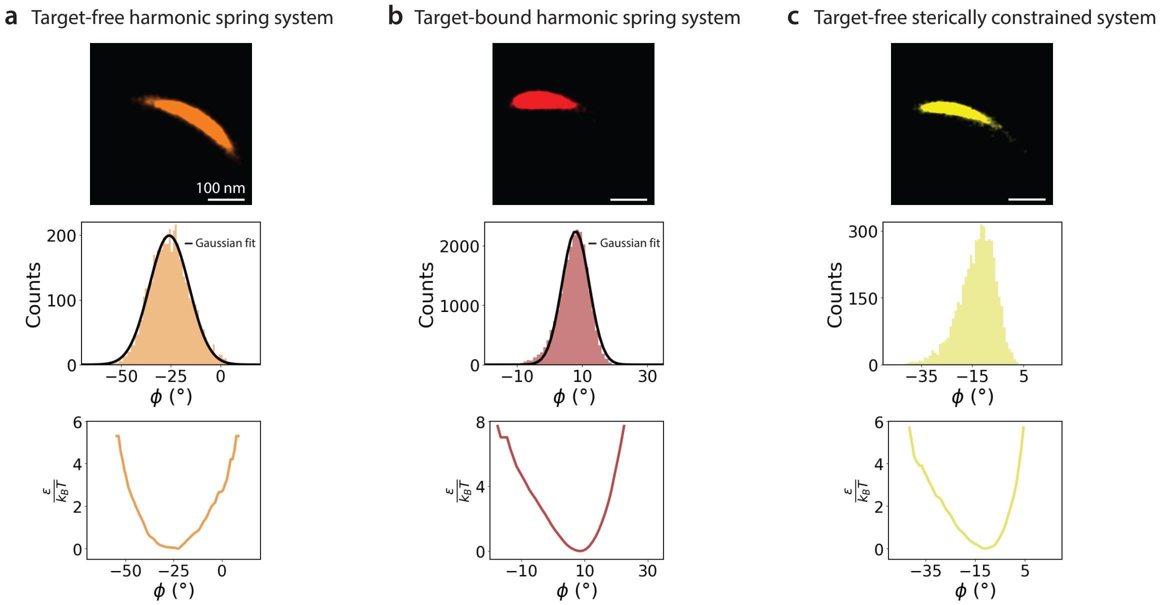

**Fig. S5**: **Mechanical characteristics of lever arm signal amplification. a-c,** Angular distributions of lever arm motion and corresponding energy landscapes. From top to bottom: Single-molecule TIRFM localization heatmaps, angular distributions of the lever arm (bin size = 1°), and the corresponding effective energy landscapes for the target-free harmonic spring system **(a)**, target-bound harmonic spring system **(b)**, and target-free sterically constrained system **(c)**. Angular distributions of the harmonic spring system under both target-free and target-bound conditions were approximated by Gaussian fits, consistent with a harmonic potential, with target-bound distribution exhibiting a small leftward shift. In contrast, the target-free sterically constrained system exhibited a left-skewed distribution, consistent with steric restriction on lever arm motion. Angular histograms were smoothed using a one-dimensional Gaussian filter ($\sigma = 1$ bin) to reduce statistical noise and normalized by their total sum to obtain the angular probability distribution, $p(\phi)$. The probability distributions were converted into effective energy landscape, $\epsilon/k_BT$, through Boltzmann inversion according to $\epsilon/k_BT = -ln(p(\phi)Z)$, where $Z$ represents the normalization factor associated with the discretized angular states. Angular bins with zero probability were excluded, and the resulting energy landscapes were shifted such that the minimum energy was set to 0.

### Steric hindrance system

The high programmability of DNA origami enables the nanomechanical amplifier's readout mechanism to be extended beyond the spring-compliant mode to a configuration in which target binding imposes a steric constraint on lever arm motion, hereafter referred to as the steric hindrance system. This is achieved by positioning recognition strands at the rear of the lever arm, on the opposite side of the hinge relative to the elastic spring system (Supplementary Fig. S6). The steric hindrance system contains 12 integration sites positioned at different distances from the hinge, denoted as peripheral, midpoint, and proximal sites in order of increasing proximity to the hinge. In this detection mode, the magnitude of the binding-induced change in lever arm motion can be tuned by varying the $Mg^{2+}$ concentration of the buffer. At low $Mg^{2+}$ concentration, the hinge remains predominantly open, limiting the transmission of the steric constraint imposed by target binding to the lever arm, resulting in a smaller displacement of the lever arm tip ($\Delta\phi$). Increasing the $Mg^{2+}$ concentration presets the hinge towards a more closed configuration, enhancing the mechanical response to the steric constraint and producing a larger angular displacement. With a single 21 nt recognition strand positioned at the proximal site, binding of a complementary 21 nt target strand produced a median $\Delta\phi$ response pooled from 106 platforms that shifted towards more negative values, from $-10.4°$ at 5 mM $Mg^{2+}$ to $-16.7°$ at 12 mM $Mg^{2+}$ and $-16.9°$ at 20 mM $Mg^{2+}$. Thus, tuning the $Mg^{2+}$ concentration provides an additional means of enhancing the angular response to target binding.

We next quantified the system's response to target strands ranging from 7 to 21 nt in two-base increments (Supplementary Fig. S7). The median response (n=77) exhibited an approximately linear dependence on target lengths, corresponding to a lever arm displacement of $\approx -1.9°$ per two base pairs increment over the measured range. Binding-induced changes in lever arm fluctuations ($\Delta\sigma$) ranged between $-0.1°$ and $-0.4°$ across target lengths, corresponding to an average reduction of approximately $3.8\%$ in angular fluctuations relative to the free strand. The increase in the magnitude of lever arm displacement with target length, without substantially altering the amplitude of lever arm fluctuations, is consistent with a steric hindrance mechanism. We then examined the effect of recognition strand position relative to the hinge. With a single 21 nt recognition strand positioned at the peripheral, midpoint or proximal site, binding of a complementary 21 nt target strand, with $\Delta\phi$ values pooled from 280 platforms, produced median responses of $-16.9°$, $-15.8°$, and $-6.6°$, respectively. Implementing a multivalent architecture by increasing the total number of recognition strands across these positions enables simultaneous binding of multiple target strands, thereby producing a cumulative steric constraint on lever arm motion. Platforms bearing six 21 nt recognition strands, with two strands positioned at each location, exhibited enhanced median $\Delta\phi$ responses upon binding of complementary target strands compared with the single-duplex readout. For a 7 nt target, median response increased in magnitude from $-2.7°$ for a single duplex to $-11.3°$ for six duplexes. At longer target lengths, lever arm displacement approached saturation, with median responses of

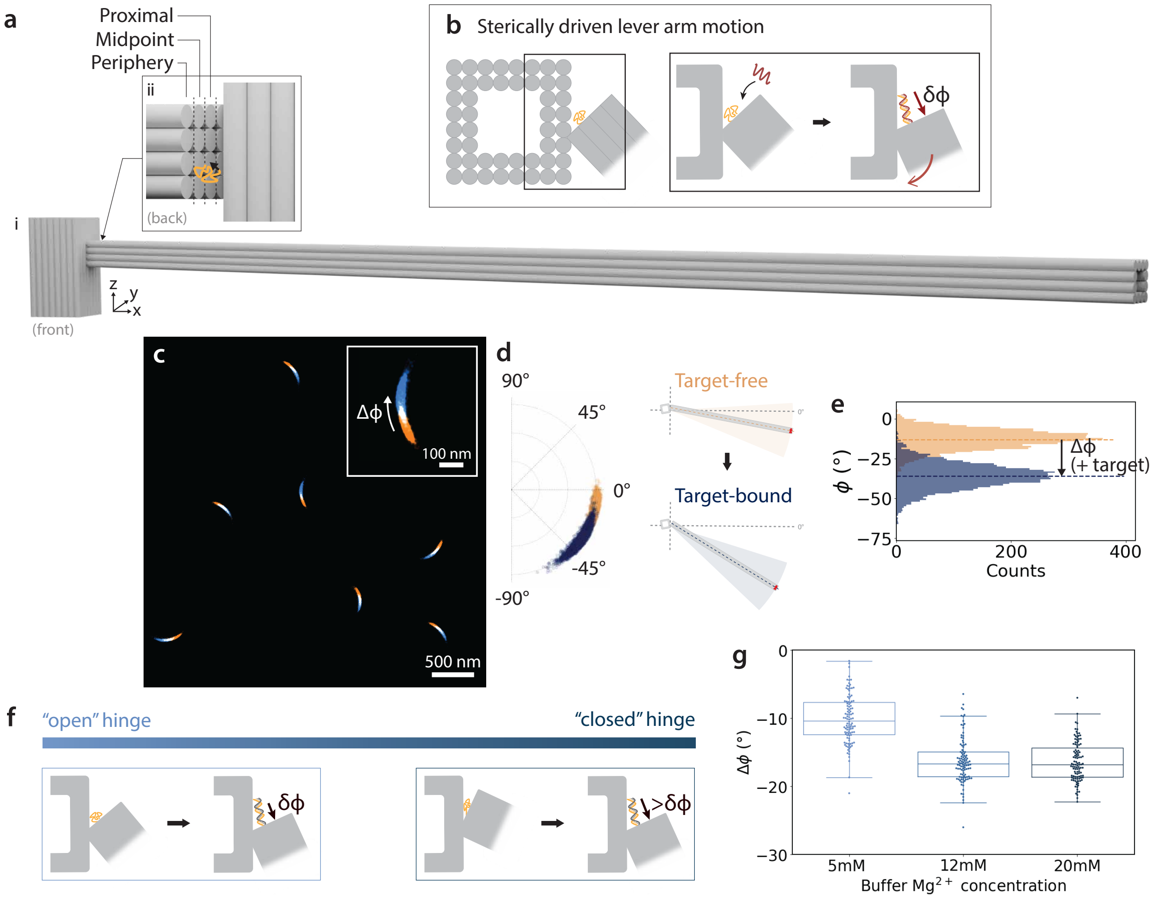

**Fig. S6**: **Steric hindrance sensing mechanism. a,** Side-view schematics of the platform viewed from the **(i)** front and **(ii)** back, with an enlarged view of the hinge-proximal terminus of the lever arm from the back (boxed). 12 recognition strand integration sites on the rear of the lever arm at three distances from the hinge—peripheral, midpoint, and proximal, ordered from furthest to closest—are indicated (each circle denotes one integration site). A schematic representation of a recognition strand integrated at the hinge-proximal site is shown (arrow indicates the integration site). **b,** Top-view schematic of the platform corresponding to the configuration shown in **a(ii)**. Hybridization of the target strand to the recognition strand forms a rigid duplex that induces steric hindrance at the hinge, driving a clockwise rotation of the lever arm about the hinge. **c,** Overlaid TIRFM localization heatmaps showing lever arm motion before (beige) and after 21 nt target strand binding (blue), with an enlarged view displaying the localization cluster corresponding to a single platform (angular positions shown in d). **d,** Polar plot of the re-referenced angular positions (left), with top-view schematics illustrating lever arm motion in target-free (top right) and target-bound (bottom right) state. **e,** Corresponding angular distributions of the lever arm trajectories. **f,** Schematic illustrating the open (low $Mg^{2+}$) and closed (high $Mg^{2+}$) hinge configurations and their influence on the binding-induced hinge-angle change ($\delta\phi$). **g,** $\Delta\phi$ values pooled from 106 platforms across all buffer conditions.

$-16.8°$ for single duplex and $-21.6°$ for six duplexes at 21 nt. Across the measured range, the median response showed a weakly nonlinear dependence on target length, with an approximately linear slope of $\approx -1.6°$ per two base pairs increment.

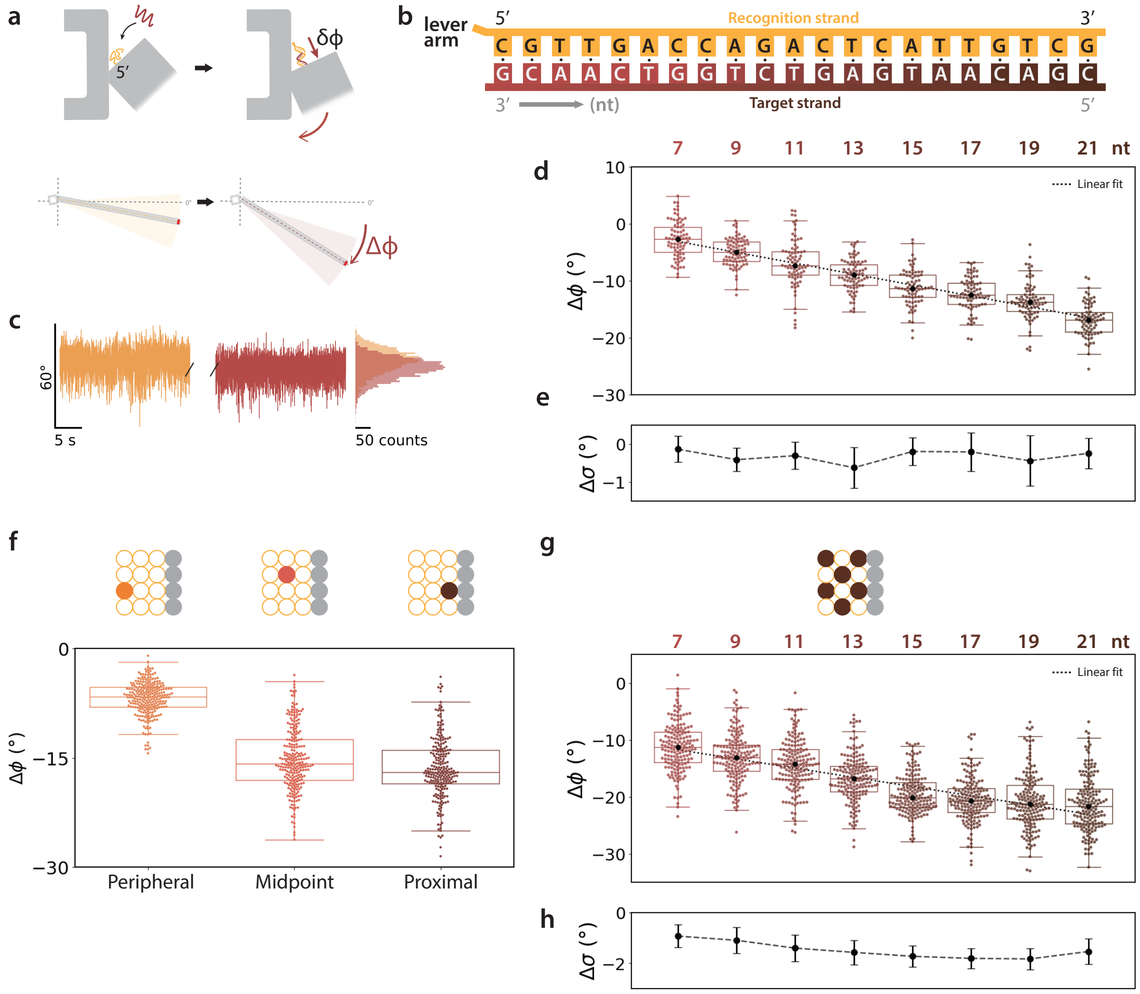

**Fig. S7**: **Angular response of sterically constrained system. a,** Top-view schematic of the platform illustrating target-strand hybridization to the recognition strand, imposing steric constraint that drives clockwise displacement of the lever arm ($\Delta\phi < 0$). **b,** Each platform contains a single 21 nt recognition strand positioned at the hinge-proximal site of the lever arm, extending from the 5′-end. Target strand lengths were increased in two-base increments towards the 3′-end of the recognition strand, ranging from 7 to 21 nt in each round of independent measurements. **c,** Representative single-lever arm time traces before (orange) and after (red) binding of a 9 nt target strand, with corresponding angular histograms. **d,** $\Delta\phi$ values pooled from 77 platforms across all measurements. **e,** Binding-induced changes in lever arm fluctuations ($\Delta\sigma$). **f,** $\Delta\phi$ values measured upon binding of a 21 nt target strand to a single recognition strand positioned at peripheral, midpoint, or proximal site relative to the lever arm hinge (exact positions depicted in the integration-site sketch above), pooled from 280 platforms. **g,** $\Delta\phi$ values measured across target lengths ranging from 7 to 21 nt in two-base increments, using platforms bearing six recognition strands, with two strands positioned at each location (peripheral, midpoint, and proximal; exact positions depicted in the integration-site sketch above), pooled from 168 platforms. **h,** Binding-induced changes in lever arm fluctuations ($\Delta\sigma$).

### Sensitivity to secondary structure in the unpaired recognition element

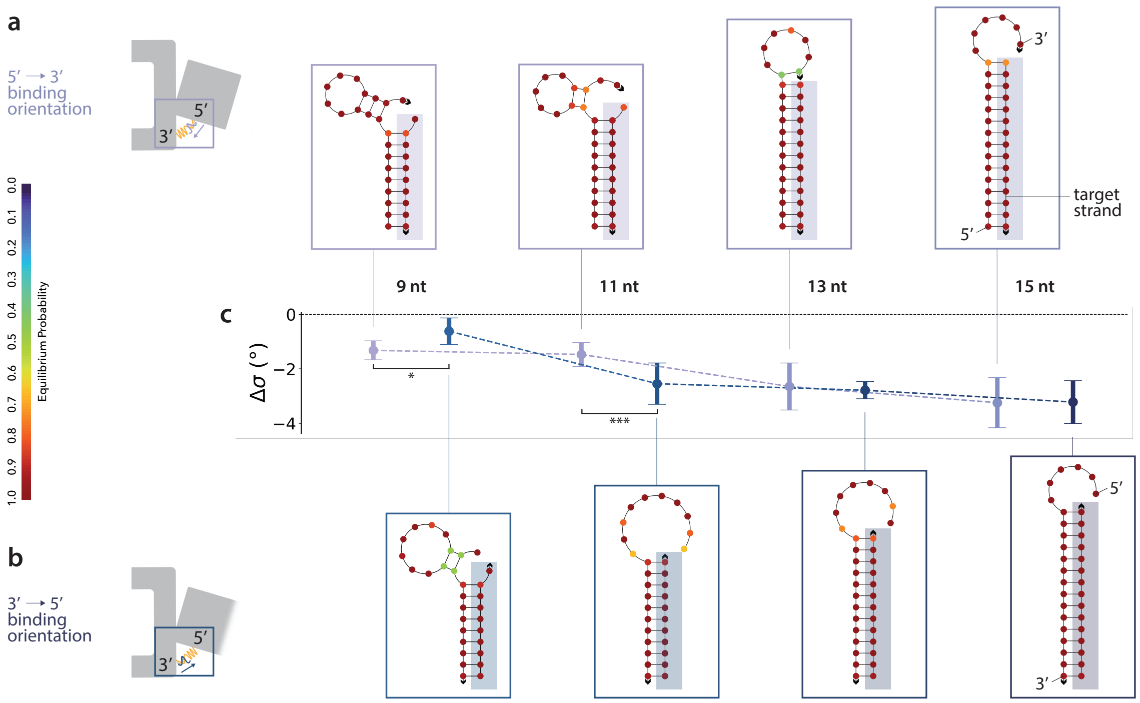

**Fig. S8**: **Angular response to secondary structure. a,b,** Top-view schematics of the platform incorporating a harmonic spring system, illustrating target-strand binding to the recognition strand in either 5'→3' **(a)** or 3'→5' **(b)** orientation, together with corresponding NUPACK-predicted equilibrium conformations of recognition–target duplexes (with the target strands highlighted using colors employed in the plots) for target lengths of 9, 11, 13, and 15 nt. **c,** Binding-induced changes in lever arm fluctuations ($\Delta\sigma$) comparing different loose-end conformations. Hairpin-containing geometries exhibited smaller reductions in angular fluctuations compared with non-hairpin geometries for 11 nt ($-1.5°$ with a hairpin vs. $-2.6°$ without a hairpin, $p = 7 \times 10^{-4}$). Statistical significance was assessed with two-sided Welch's t-test with $\alpha = 0.05$ Holm adjustment; *$P < 0.05$, ***$P < 0.001$.

### Parallelized monitoring of strand–exchange dynamics

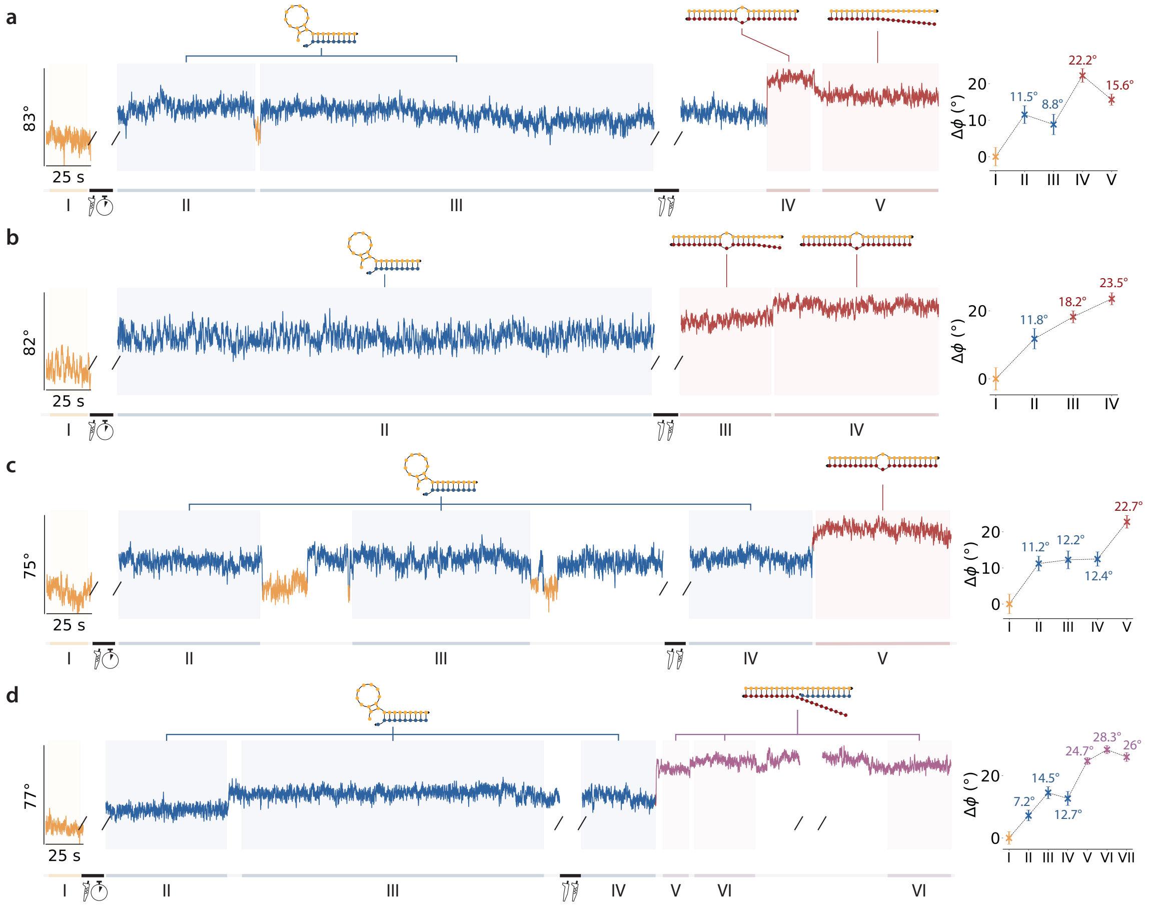

**Fig. S9**: **Parallel single-molecule monitoring of strand-exchange dynamics. a-d,** Left: Selected denoised single-molecule traces from individual platforms acquired within the same field of view ($N_{total} = 183$), capturing strand-exchange dynamics of 9 nt and 21 nt single-base-mismatch target strands interacting with a single 21 nt recognition strand. Putative strand interactions inferred from angular displacement ($\Delta\phi$) analysis are illustrated in the schematics above each trace. Right: Segment-specific $\Delta\phi$ values referenced to the free strand baseline.

### Dopamine-binding aptamer dynamics

Kinetic heterogeneity of the dopamine-binding aptamer transitions between elongated and contracted states was quantified by model comparison using the Bayesian information criterion (BIC) (Supplementary Fig. S10). The double-exponential model was favored over the single-exponential model for both states at all dopamine concentrations ($\Delta\mathrm{BIC} = \mathrm{BIC}_{\mathrm{single}} - \mathrm{BIC}_{\mathrm{double}} > 100$). However, the improvement provided by the double-exponential model was only marginal for the elongated state across all concentrations and for the contracted state in the absence of dopamine. For the elongated state, the fast component accounted for 92–96% of the fitted amplitude across 10–100 µM dopamine. The forward transition is therefore governed primarily by the fast component, likely reflecting G-quadruplex folding, with dopamine binding contributing, at most, a minor kinetically distinct pathway. In contrast, the contracted state at 10, 50, and 100 µM dopamine exhibited substantially larger $\Delta\mathrm{BIC} > 8000$ values, indicating decisive statistical support for kinetically heterogeneous dwell-time populations.

We then examined the lever arm angular response to determine whether G-quadruplex-induced conformational changes could be distinguished from dopamine-binding events (Supplementary Fig. S12). Based on the transition rates $k_{\mathrm{CE,fast}}$ and $k_{\mathrm{CE,slow}}$ associated with the dissociation pathway (C$\rightarrow$E), angular traces were classified into fast dissociation events, likely corresponding to unfolding of the intrinsic G-quadruplex conformation, and slow dissociation events, likely corresponding to unfolding of a dopamine-stabilized G-quadruplex conformation. Lever arm displacements pooled from 91 structures exhibited Gaussian-fitted distributions with an approximately constant peak for the slow pathway ($\mu_{slow} \approx 18.5°$) across all dopamine concentrations. In contrast, the fast pathway exhibited a modest increase in the fitted peak, from $\mu_{fast} \approx 19.1°$ at 10 µM to $\mu_{fast} \approx 19.7°$ at 50 µM and $\mu_{fast} \approx 20.3°$ at 100 µM. Accordingly, the difference between the two pathways ($\mu_{fast} - \mu_{slow}$) increased modestly from $0.6°$ at 10 µM to $1.2°$ at 50 µM and $1.8°$ at 100 µM. As the pathway-dependent shift remained small and the displacement distributions closely overlapped, this suggests that the same structural transition underlies both processes. This interpretation is consistent with previous reports showing that ligand association and dissociation occur on substantially shorter timescales than the conformational change [2]

The overall mean lever arm positions ($\bar{\phi}$) for the elongated and contracted states revealed an approximately invariant lever arm displacement ($\Delta\phi$) across dopamine concentrations: $17.5°$ at 0 µM, $18.6°$ at 10 µM, $18.1°$ at 50 µM, and $18.2°$ at 100 µM. Increasing dopamine concentration primarily redistributed the conformational equilibrium toward the contracted state. The contracted-state occupancy increased from 15.6$\pm$1.1$\%$ at 0µM to 69.7$\pm$1.4$\%$ at 100µM dopamine, accompanied by a corresponding reduction of the elongated-state occupancy from 84.4$\pm$1.1$\%$ to 30.3$\pm$1.4$\%$.

We further observed substantial inter-platform heterogeneity in both the number of fluctuations from the elongated state and the median elongated-state lifetime. A two-dimensional Gaussian

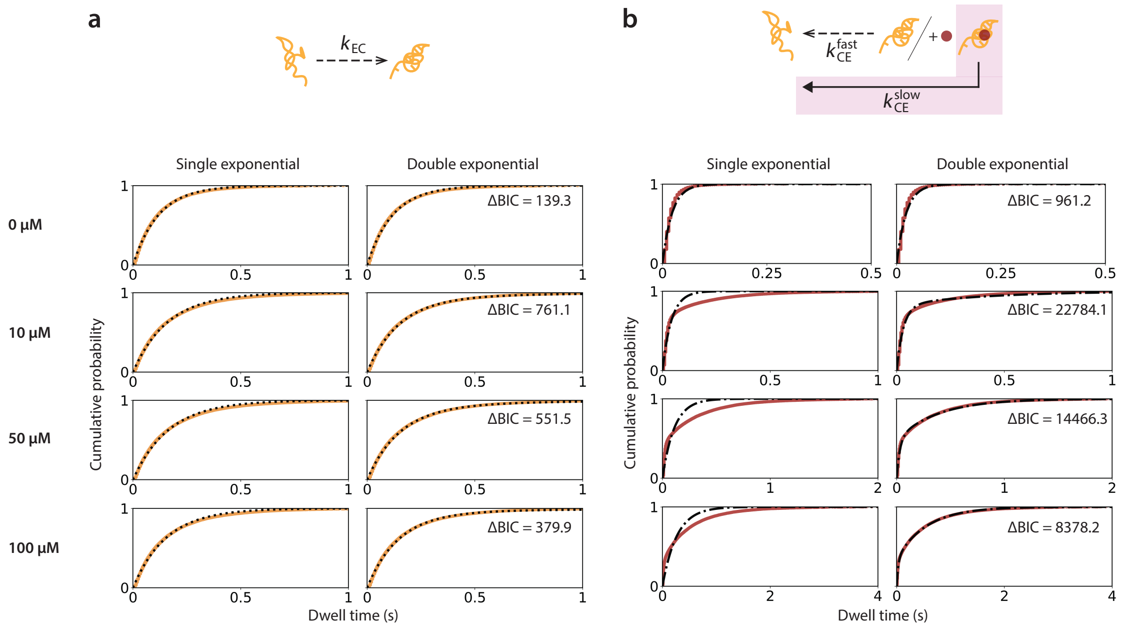

**Fig. S10**: **Comparison of single- and double-exponential fits to dopamine-binding kinetics. a,b,** Top: Schematics illustrating the transitions of dopamine-binding aptamer from elongated to contracted state **(a)** and vice versa **(b)**. Bottom: Empirical cumulative distribution functions (ECDFs) of HMM-extracted dwell times from 91 platforms for the elongated state (**a**; corresponding to $k_{\mathrm{EC}}$) and contracted state (**b**; corresponding to $k_{\mathrm{CE}}$) at increasing dopamine concentrations (0, 10, 50, and 100 μM; data underlying Fig. 4). For each concentration, ECDFs were fitted with single- and double-exponential cumulative distribution models (black lines).

mixture model (GMM) with BIC-based model selection identified three subpopulations (Supplementary Fig. S13): a fast-switching population characterized by frequent transitions and short median elongated-state lifetimes ($\approx$ 11%, n=10), a slow-switching population exhibiting infrequent transitions and prolonged lifetimes ($\approx$ 18%, n=16), and a dominant intermediate population exhibiting mixed behavior ($\approx$ 71%, n=65). Population-resolved kinetic analysis revealed heterogeneity in apo-state transition rates that was otherwise masked by ensemble averaging. Specifically, $k_{\mathrm{EC}}$ values were $13.3\ \mathrm{s}^{-1}$, $8.2\ \mathrm{s}^{-1}$, and $4.8\ \mathrm{s}^{-1}$ for the fast-switching, dominant, and slow-switching populations, respectively, whereas $k_{\mathrm{CE}}$ values were $36.8\ \mathrm{s}^{-1}$, $44.9\ \mathrm{s}^{-1}$, and $47.3\ \mathrm{s}^{-1}$.

In the presence of dopamine, a second dissociation pathway ($k_{\mathrm{CE,slow}}$) emerged and remained largely invariant across dopamine concentrations and subpopulations, ranging from $1.3\,\mathrm{s}^{-1}$ to $2.7\,\mathrm{s}^{-1}$. In the fast-switching population, the faster dissociation pathway ($k_{\mathrm{CE,fast}}$) decreased modestly with increasing dopamine concentration, from $30.7\ \mathrm{s}^{-1}$ at 10 μM to $27.1\ \mathrm{s}^{-1}$ at 50 μM and $24.7\ \mathrm{s}^{-1}$ at 100 μM. This observation suggests that dopamine may selectively stabilize a pre-existing G-quadruplex conformation, thereby shifting the conformational equilibrium toward the contracted state. In contrast, $k_{\mathrm{CE,fast}}$ remained largely independent of dopamine concentration in the dominant and slow-switching populations, with average values of $33.7\ \mathrm{s}^{-1}$ and $39\ \mathrm{s}^{-1}$, respectively. The folding transition rate ($k_{\mathrm{EC}}$) remained largely concentration-independent across all subpopulations, with average values of $11.4\ \mathrm{s}^{-1}$, $5.8\ \mathrm{s}^{-1}$, and $4.5\ \mathrm{s}^{-1}$ for the fast-switching, dominant, and slow-switching populations, respectively.

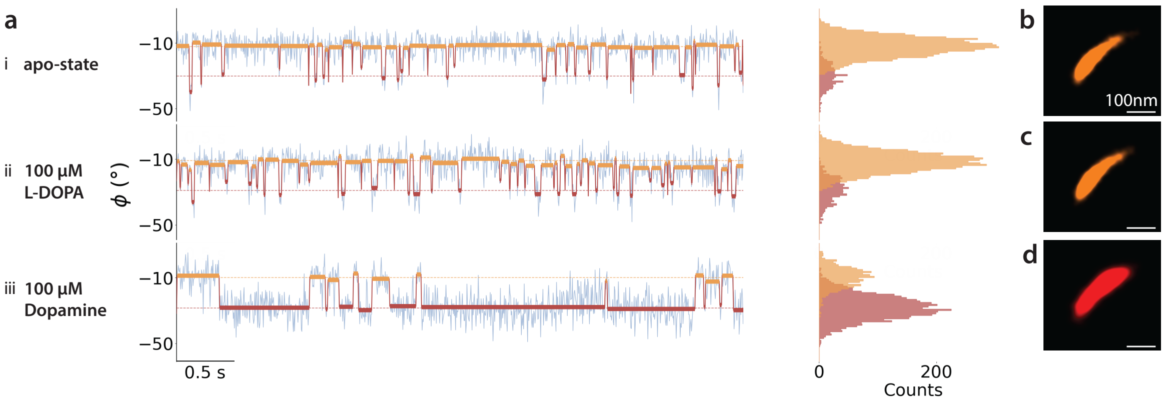

**Fig. S11**: **Target specificity of the platform. a,** Left: Representative single-molecule angular time traces of the same platform (blue, first 5 s excerpt) showing transitions between the elongated (orange) and contracted (red) aptamer states under **(i)** apo-state conditions, **(ii)** in the presence of 100 µM L-DOPA, or **(iii)** 100 µM dopamine. Right: Corresponding HMM-derived angular histograms of the elongated and contracted states (first 25 s excerpt). **b-d,** TIRFM localization heatmaps showing lever arm motion during the first 25 s under apo-state conditions **(b)**, 100 µM L-DOPA **(c)**, and 100 µM dopamine **(d)**.

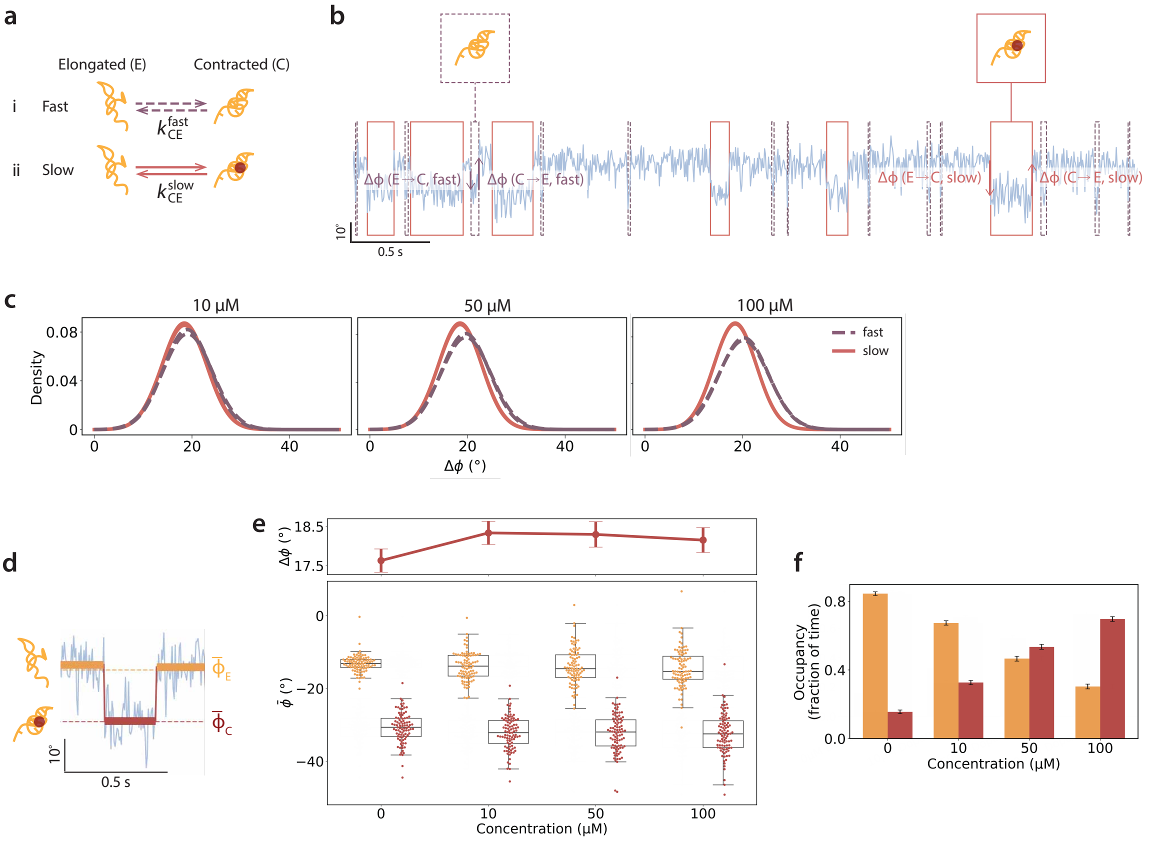

**Fig. S12**: **Angular response to dopamine binding. a,** Top: Schematics illustrating the transition of dopamine-binding aptamer from an elongated (E) conformation to a G-quadruplex-induced contracted (C) conformation stabilized upon dopamine binding. Distinct transition rates ($k_{\mathrm{CE,fast}}$ and $k_{\mathrm{CE,slow}}$) were observed for the dissociation pathway. **b,** Single-molecule angular time trace (blue, 5 s excerpt) showing contracted-state intervals associated with fast (purple brackets) and slow (carmine brackets) dissociation pathways. **c,** Lever arm displacements ($\Delta\phi$) pooled from 91 platforms for forward (E→C) and backward (C→E) transitions at 10, 50, and 100 µM dopamine concentrations. Plotted curves represent the Gaussian fits to the angular distributions, and the means ($\mu$) reported correspond to the peaks. **d,** Enlarged view of single-molecule angular time traces showing the interconversion between aptamer states. The thick lines indicate the mean lever arm position of each state dwell, and the dotted lines indicate the per-platform mean lever arm positions of the elongated ($\bar{\phi}_E$, orange) and contracted ($\bar{\phi}_C$, red) state. **e,** Mean lever arm positions of both states across dopamine concentrations (bottom, n=91) and corresponding lever arm displacements (top). **f,** State occupancy, quantified as the fraction of time aptamers occupied each conformational state, shown as mean±SEM.

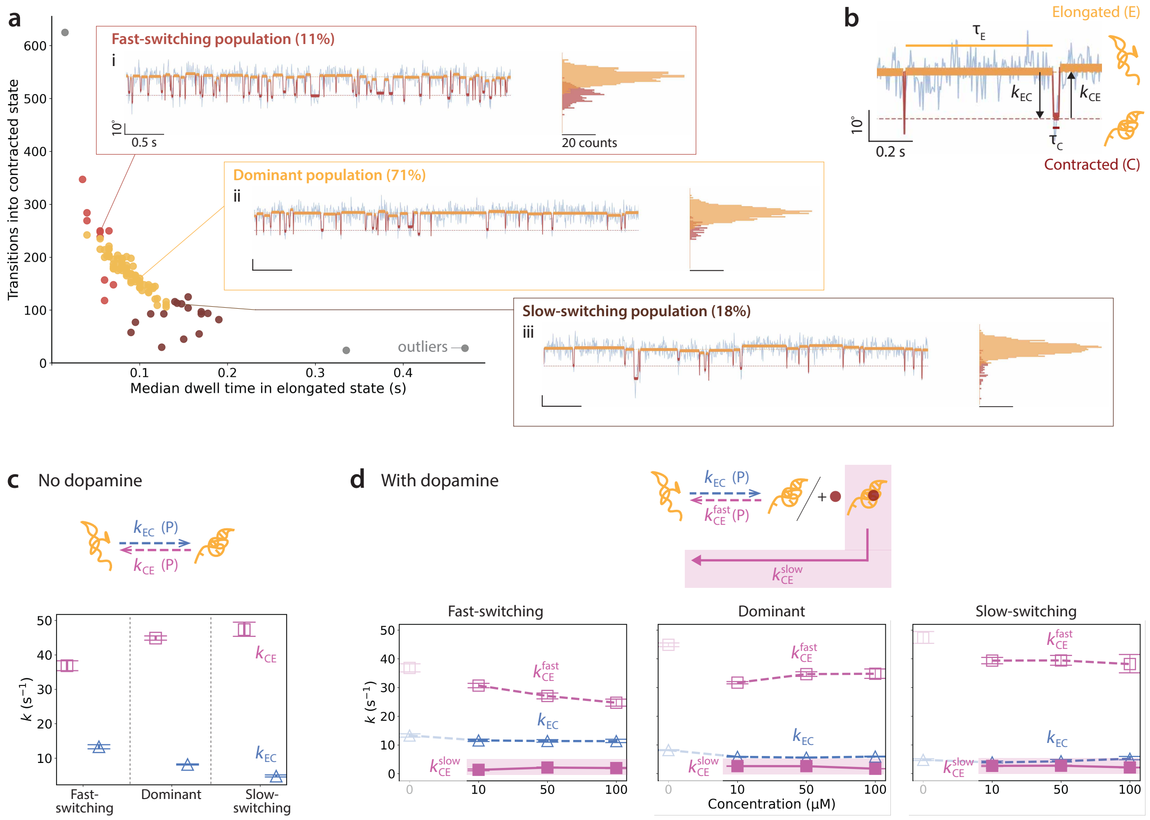

**Fig. S13**: **Population-based analysis of dopamine-binding kinetics. a,** Two-dimensional GMM analysis of median elongated state lifetime and number of transitions into the contracted state for the 25 s apo-state measurement. BIC-based model selection identified three sub-populations with distinct kinetic behaviors: **(i)** fast-switching (representative single-molecule angular time trace shown), **(ii)** slow-switching, and **(iii)** dominant population. **b,** Enlarged view of a single-molecule angular time trace showing transition between the elongated (E) and contracted (C) states. Transition rates were obtained by fitting the ECDFs of the elongated state dwell times ($\tau_E$, corresponding to $k_{\mathrm{EC}}$) and contracted state dwell times ($\tau_C$, corresponding to $k_{\mathrm{CE}}$). **c,** Apo-state transition rates of each sub-population (schematic illustrating the state transitions shown above). **d,** Transition rates of each sub-population across 10, 50, and 100 μM dopamine (schematic illustrating the state transitions shown above).

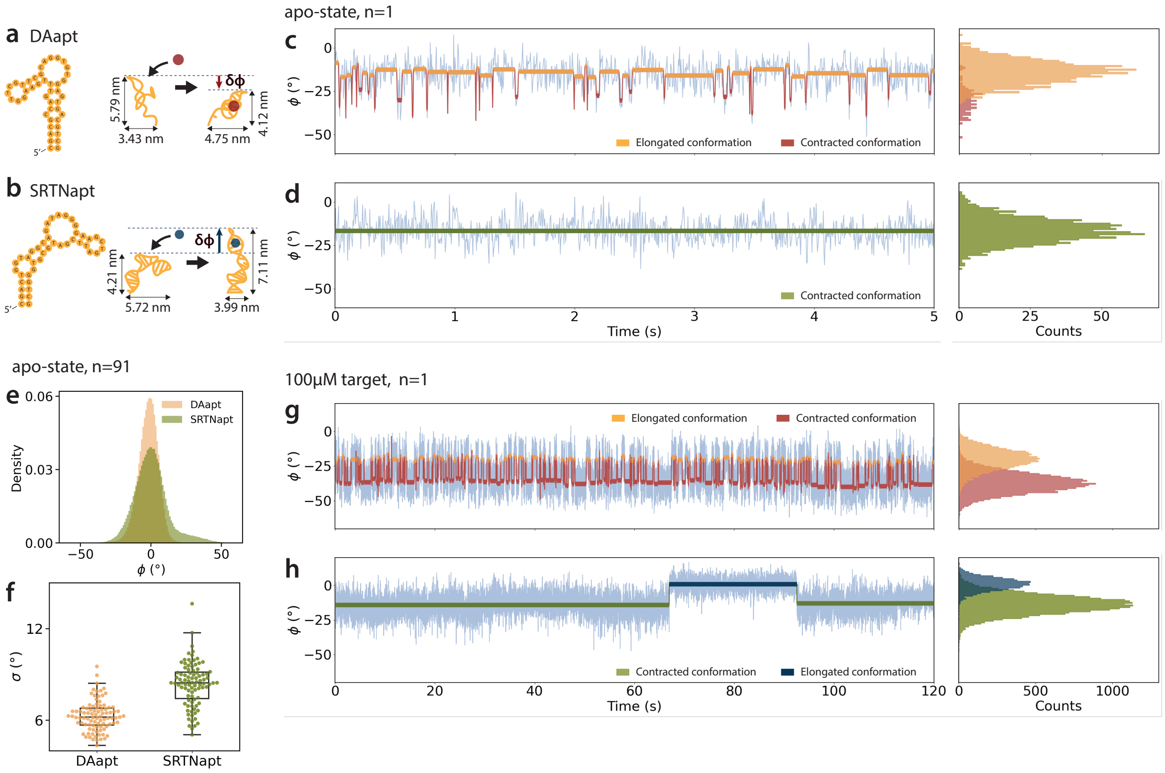

**Fig. S14**: **Dopamine-binding versus serotonin-binding aptamer. a,b** Schematics illustrating the dopamine-binding (DAapt, **a**) and serotonin-binding (SRTNapt, **b**) aptamers reported by Nakatsuka et al. [3]. Both aptamers adopt G-quadruplex conformations, inducing a conformational change that results in contraction of DAapt or elongation of SRTNapt, respectively [4]. **c,** Representative single-molecule angular time trace (blue, first 5 s excerpt) of an apo-state DAapt showing interconversion between the elongated conformation (orange) and G-quadruplex-induced contracted conformation (red), with corresponding angular histograms. **d,** Representative single-molecule angular time trace of an apo-state SRTNapt showing a single dominant state (green) without discernible interconversion. **e,** Lever arm angular positions pooled from 91 structures revealed a broader angular distribution for the unfolded SRTNapt state compared with the unfolded DAapt state. **f,** Lever arm fluctuations ($\sigma$) further revealed more constrained motion for the DAapt-integrated platform compared with the SRTNapt-integrated platform (median $\sigma_{DAapt} = 6.2°$, $\sigma_{SRTNapt} = 8.5°$). This difference may arise from increased stem fraying in SRTNapt, which contains a shorter 4-bp stem designed to stabilize the adjacent G-quadruplex loop, thereby reducing structural stability compared with DAapt. **g,** Single-molecule angular time trace showing HMM-fitted elongated (orange) and contracted (red) aptamer states in 100 µM dopamine, with corresponding angular histograms. **h,** Single-molecule angular time trace showing HMM-fitted contracted (green) and elongated (blue) aptamer states in 100 µM serotonin, with corresponding angular histograms. Unlike the frequent interconversion between states observed for DAapt at saturating ligand concentrations, SRTNapt exhibited stochastic ligand-induced transitions, likely reflecting reduced availability of the ligand-binding G-quadruplex conformation due to reduced structural stability. The markedly extended bound-state lifetime of SRTNapt suggests different kinetic characteristics compared with DAapt.

### Nanomechanical amplifier folding sequences

Staples included in the one-pot folding reaction for each structural variant are highlighted in green. The structural variants denoted as **spring** and **steric** correspond to the elastic spring system and steric hindrance system, respectively (Supplementary Table 1). **1x** and **6x** indicate the number of recognition strands incorporated at the rear of the lever arm, adjacent to the hinge region, for target detection. The terms **1x-Proximal**, **1x-Midpoint** and **1x-Peripheral** denote the respective incorporation sites of the recognition strand (Supplementary Table 3). For the different recognition strands used for target measurements (Supplementary Tables 2 and 3), **L1-21nt** indicates the 21 nt single-stranded DNA (ssDNA) recognition strand used for strand hybridization experiments, whereas **P1** indicates the ssDNA recognition strand used for DNA-PAINT experiments. **DAapt** and **SRTNapt** indicate the dopamine-binding and serotonin-binding aptamers, respectively. Modified strands are marked using the same colors as indicated in Supplementary Fig. S1. ATTO655-labeled strands used for lever arm localization tracking are LNA-modified strands (Supplementary Table 4). Lever arm staples and platform base staples were folded separately using the p8064 scaffold.

### Table S1. Platform base staple.

| Oligo Name | Sequence (5' to 3') | Spring | Steric-1x | Steric-6x |
|---|---|---|---|---|
| 0[107]1[107] | TTTT TCTTTGACCCCCAGCGTACACTAAAACACTCA TTTT | | | |
| 0[47]45[47] | ATCATCGCTTTCGTCTAGTTAAACTCCCGTAA | | | |
| 0[79]46[72] | CAAGCGCGGCACATCCCAGCAGCA | | | |
| 1[15]1[39] | TTTT CCGCGACCTGCTCCATGTTA | | | |
| 1[40]4[40] | CTTAGCCGAGGACAGATCTTGACATAAGGCTT | | | |
| 1[64]0[80] | GTCAATCATAAGAGGCAAAAGAAATTATAC | | | |
| 2[102]3[79] | TTTT GAAGGCACCAACCTAAAACGAAAGGGTCAGGTAA | | | |
| 2[55]0[48] | TTTGAAAGGAACGAGGAGATTTGT | | | |
| 3[48]44[48] | AGAACCGGAAGGGTAACGTCGCTGACTTAAAT | | | |
| 3[80]44[80] | AATACGTGTCTGGTTCATAACAGCTTAC | | | |
| 3[9]0[15] | TTTT GGCTGGCTGACCTTCATCAATTGTGTCGAAAT TTTT | | | |
| 4[107]5[107] | TTTT TTTTTCATGAGGAAGTTTTGAGGACTAAAGAC TTTT | | | |
| 4[39]9[39] | GCCCTGATAACGGACCAGTCAGAGCAAC | | | |
| 4[79]9[71] | AAACGACGTAACAAAGTTCATCAGCCTTATGCACGACGAT | | | |
| 5[40]8[40] | TGAGATGGTCATTATAACAACATTTGCAGATA | | | |
| 5[70]8[72] | GTGAAGGCTAGCAACTTATTGCGTCAGGGATAGA | | | |
| 6[102]4[80] | TTTT AGCATCGGAACGAGGGTACAGAGGCTTCCATT | | | |
| 6[55]4[48] | GAACTGGCTTTAATTTTCAGTGAA | | | |
| 7[56]2[56] | TAGAAAGACTGCTCATCAACTTTATGACCAAC | | | |
| 7[9]4[15] | TTTT CTACGTTAATAAAACGAACCGAGAAACACCAG TTTT | | | |
| 8[107]9[107] | TTTT AGGGAGTTAAAGGCCGGGTCGCTGAGGCTTGC TTTT | | | |
| 8[39]13[39] | CATAACGAATACTGTTGCCAGATAGTCA | | | |
| 8[71]13[71] | TTTAGGAAAATCCCCCCAAAATAGAGATTAAG | | | |
| 8[87]7[102] | CTTTTGCCCCTCAGCAGCGAAAGAC TTTT | | | |
| 9[40]12[40] | ACTATCATAAGAAGTTCGGAATCGAATGACCA | | | |
| 9[72]12[72] | AAAACCGAGCATAAACTCATACCGCTTGAAAT | | | |
| 10[102]9[87] | TTTT ACAACAACCATCGCCCACTATATTC | | | |
| 10[55]7[55] | TTTTGCAAAACCCTCGTCAACTAAATTACAGG TT/3ATTO488N/ | | | |
| 11[56]6[56] | ATTCATTGTACCACATTTTACCAGGATTTTAA | | | |
| 11[9]8[15] | TTTT TTTAGACTGGATAGCGTCCCCAAAAGGAATTA TTTT | | | |
| 12[107]13[107] | TTTT TTCGAGGTGAATTTCTGGTTTATCAGCTTGCT TTTT | | | |
| 12[71]14[56] | GCTTTAAAGCGAATAACCCGAAAGACTTCAAA | | | |
| 13[40]15[47] | GAAGCAAATTTAATTCGAGTACCT | | | |
| 13[72]11[102] | AGGTTTAATTGTATCTAAACAGATAGTTGCGCCGACAATG TTTT | | | |
| 14[102]15[79] | TTTT AAAAAAAGGCTCCAAAAGGAGCCAAGTAATTTTT | | | |
| 14[55]11[55] | TATCGCGTGCGGATTGAAAACGAGTCATAAAT | | | |
| 15[48]10[56] | TTAATTGCTCCTTATTCAGTTCAGCATCAAAACGAGAGGC | | | |
| 15[80]18[80] | TCACGTTGAGAATACTGTATGGATCTAA | | | |
| 15[9]12[15] | TTTT GCAAACTCCAACAGGTCAGAAAATCAGGTCTT TTTT | | | |
| 16[110]15[102] | TTTT TCAACAGTTTCAGCGGAGTGAAAATCTCCA TTTT | | | |
| 16[65]17[71] | AAGGATTGATAAGAGGTCCATGTTTTAAATAACGTTAGTAAA | | | |
| 17[17]12[32] | TTTT TTGCTGAATATAATGCTGTAAATCA | | | |
| 17[40]19[55] | TAGCTCAAATTTTTGCGTAGATTTAGTGTCAC | | | |
| 17[72]19[79] | TGAATTTTGAAAGGAAGCCTGTAG | | | |
| 18[104]17[110] | TTTT TAGTTAGCGTAACGGATTTTGCTAAACAACTT TTTT | | | |
| 18[47]23[47] | CGGTGTCTTATTTTCAACCATTAGCAAAGAAT | | | |

| Oligo Name | Sequence (5' to 3') | Spring | Steric-1x | Steric-6x |
|---|---|---|---|---|
| 18[79]23[79] | AGTTTTGTTAAATCAAATACAAATGCAAGCGG | | | |
| 19[32]16[17] | CGAACGAGGATGGCTTAGAGCTTAA TTTT | | | |
| 19[56]18[48] | CAGTACAACAGTGCAACTAAAGTA | | | |
| 19[80]22[80] | CATTCCAAGAGTCCAAGAATATGTTTGA | | | |
| 20[110]19[104] | TTTT GTTGTTCCAGTTTGGAACACAGACAGCCCTCA TTTT | | | |
| 20[63]21[55] | GTATAAGAGTTTCTTGTTTGGGGC | | | |
| 21[17]18[32] | TTTT CAATAACCTGTTTAGCTAGGAAGTT | | | |
| 21[56]16[66] | GCGAGAAAATCCCTTACGTCTTTCACTACAACCAACTA | | | |
| 22[104]21[110] | TTTT CAGGCGAAAATCCGCCCGAGATAGGGTTGAGT TTTT | | | |
| 22[47]25[47] | ATCAATTCGGTAAAGAGAGCATAAGAAGCCTT | | | |
| 22[79]25[79] | TGGTGGTTCGCTTTCCCCTTCACCGGTGGTTT | | | |
| 23[48]22[48] | TAGCAAAATTAAGAGAGGCCTGAAAAGGTGGC | | | |
| 23[80]27[87] | TCCACGCTGATTGCAGTCGGG | | | |
| 24[102]23[104] | TTTT CGGGCAACAGCTGGTTTGCCCCAG TTTT | | | |
| 24[31]22[12] | TCGGTTGTATCATACAAGTAGTAGCATTAACA TTTT | | | |
| 25[80]25[102] | TTCTTTTCACCAGTGAGA TTTT | | | |
| 26[107]27[107] | TTTT CTGCATTAATGAATCGAAACCTGTCGTGCCAG TTTT | | | |
| 26[39]24[32] | ACCCTCATGCGGGAAGCTAAA | | | |
| 26[71]27[63] | GAGGCGGTTTGCGAAGGGTGAGAA | | | |
| 26[87]29[102] | GCCAACGGGTGCCTAATGAGTGAGC TTTT | | | |
| 27[15]28[9] | TTTT CAATGCCTGAGTACGTTCTAGCTGATAAATTA TTTT | | | |
| 27[36]20[17] | GTATACTAATGGCAAGGATACATTTCGCAAATGGT TTTT | | | |
| 27[64]20[64] | AGGCCGCCCCGAAATCGAGTTGCATCTTACCA | | | |
| 28[102]31[87] | TTTT TAACTCACATTAATTGCGTGTGAAA | | | |
| 28[55]33[55] | TCACCATCGATAATCAGCAAACAATTTTAACC | | | |
| 29[56]28[56] | ATCAGGTCATTGCGTAACTGGAGACAGTCAAA | | | |
| 30[107]31[107] | TTTT ATTCCACACAACATACTTGTTATCCGCTCACA TTTT | | | |
| 30[39]27[35] | ATGAACGTTTTTGAATTCAACATGT | | | |
| 30[73]31[73] | AAAGTCTGAGAGTCTGGAGAAAAGCCCCAAAGCTGT | | | |
| 30[87]33[102] | GAGCCGGGTTGAGGATCCCCGGGTA TTTT | | | |
| 31[15]32[9] | TTTT TAGCATGTCAATCAACGTTAATATTTTGTTAA TTTT | | | |
| 31[40]26[40] | CCCCGGTTAATATGATGAGATCTATTTTTAGA | | | |
| 31[74]26[72] | TTCCTGTTGCGCTCAAGCCTGGCGCGGGGA | | | |
| 32[102]35[87] | TTTT CCGAGCTCGAATTCGTAATTCACGG | | | |
| 32[55]37[55] | TATAAGCAATTCTCCGGCGTCTGGCGGCCTCA | | | |
| 33[56]32[56] | AATAGGAACGCCAGCCATAAACAGGAAGATTG | | | |
| 34[107]35[107] | TTTT TGCCAGCACGCGTGCCTCATACCGGGGGTTTC TTTT | | | |
| 34[39]31[39] | TAGCCAGTTAAATCAATTGTAATATGTA | | | |
| 34[74]35[74] | CCGTGATCAAAAATAATTCTGGGAACAAACGGGGCGGG | | | |
| 34[87]37[102] | TGTTCTTAGTGTCACTGCGCGCCTG TTTT | | | |
| 35[15]36[9] | TTTT TAAATGTGAGCGAGATGGGCGCATCGTAACCG TTTT | | | |
| 35[40]30[40] | CCCGTCGGAATATTTAAGCTCATTGAGAATCG | | | |
| 35[75]30[74] | CCGTTTCATGGTCTCCTCACAAAGCAT | | | |
| 36[102]41[104] | TTTT TGCACTCTGTGTAAAGGTTTCTTT TTTT | | | |
| 36[63]43[63] | TGACCGTACGTTGTAACCTCTTCGAGACGCAG | | | |
| 37[56]36[64] | GGAAGATCGCACTCGATGCCGGAT | | | |

| Oligo Name | Sequence (5' to 3') | Spring | Steric-1x | Steric-6x |
|---|---|---|---|---|
| 38[107]39[107] | TTTT TGCAGCCAGCGGTGCCCAGATGCCGGGTTACC TTTT | | | |
| 38[39]35[39] | GCCGGAAGGACGACTGGTGTAGTAACAA | | | |
| 38[74]39[74] | ATCAGACCAGCCAGCTTTCGCGATCGGTGCCAGCAAAT | | | |
| 39[15]40[32] | TTTT CCATTCGCCATTCAGGCTGCCGAAAGG | | | |
| 39[75]34[75] | CGTTAGGCCAGAATCCAGCGCCGCGT | | | |
| 40[104]43[110] | TTTT GCTCGTCATAAACCAGGCGCTTTCGCACTCAA TTTT | | | |
| 40[47]34[40] | CCAGCTGGGCAACTGTGGTCACGTGACAGTATCCTTCCTG | | | |
| 40[79]41[79] | ACACTGGTGTGTTGGGAACGCCCTGCGGCTGG | | | |
| 41[32]42[17] | CGCCAGGTTGTGAGAGATAGACTTT TTTT | | | |
| 41[48]38[40] | AGTCACGAATGGGATATGGGAAGGCGGCACCGCTTCTGGT | | | |
| 41[80]38[75] | TAATGGGGTGCTGCACGGCATGGTGCCCCCTGC | | | |
| 42[110]45[104] | TTTT TCCGCCGGGCGCGGTTGCGCGTCGGTGGTGCC TTTT | | | |
| 42[71]47[63] | CTGTTGACGGCCAGTGGCACAGGCCGGATCAAGCAGCCTC | | | |
| 43[17]44[32] | TTTT CTCCGTGGTGAAGGGATAATTTGCC | | | |
| 43[64]42[72] | AAACAGCATCAGCGGGCCGGGTCA | | | |
| 44[104]47[110] | TTTT ATCCCACGCAACCGGAACGTGCCGGACTTGTA TTTT | | | |
| 44[47]41[47] | TTCTGCTCGCTCTCACCCCGGAATGTTTTCCC | | | |
| 44[79]45[79] | GGCTGGAGGTGTCCAGGGCAAGAATGCCAACG | | | |
| 45[32]46[17] | AAAAAAAGATGCTGATTGCCGTTCC TTTT | | | |
| 45[48]40[48] | AAAAAGCCCCAAGCTTGGAAAAAGCTATTACG | | | |
| 45[80]40[80] | GCAGCACGTATGAGGTCATTGATCCCTT | | | |
| 46[110]3[102] | TTTT GAACGTCAGCGTGGTGCTGAATGCCACTAC TTTT | | | |
| 46[71]1[63] | ACCGCCTTTAGTGATGATATTCATTACAACGGCGCAGACG | | | |
| 47[17]5[39] | TTTT GGCAAACGCGGTCCGTTTCTGATAAAGAGTAATGAACGGTTGGGCT | | | |
| 47[64]5[69] | CGGCCAGAAAACAAAGTACCCAAAAACCGAACATCATT | | | |
| 48[239]60[232] | CACCGGATAACAACTTGCCTTACCTATTGAGTGTA | | | |
| 48[271]62[264] | GTGTGATACATTTTCGAGCCAGTAAACCGCCACCCTCAGA | | | |
| 48[303]62[296] | GACCTAAAGTACCGACAAAAGGTATACTCAGGAGGTTTAG | | | |
| 48[335]62[328] | TTCAAATACGACGACAATAAACATAAGTATAGCCCGG | | | |
| 48[367]62[360] | CAATCGCAGCAGAACGCGCCTGTTACTTGCGGGAGGTTTT | | | |
| 48[399]62[392] | ATATAACTAGTCCTGAACAAGAAAAAGAACGCGAGGCGTT | | | |
| 48[431]62[424] | TACCTTTAATTTACGAGCATGTAATCAGATATAGAAG | | | |
| 49[232]55[248] | TCTATCAGCGCGTACACCAGTCACAC TTTT | | | |
| 49[264]51[263] | ATCACCCAGCTAGGGCGCTGGCAAAGAGCGGG | | | |
| 49[296]51[295] | TGCCGTAAAAAGGAAGGGAAGAAAGGGATTTT | | | |
| 49[328]48[336] | GAGCCCCCGATTAGAAAAACTTT | | | |
| 49[360]51[359] | ATCAAGAAATTATTCATTTCAATTACATTATC | | | |
| 49[392]51[391] | ATTTCATTAATACCAAGTTACAAACAGAAGGA | | | |
| 49[424]51[423] | AGTACATGAAACAATAACGGATTTATCAGA | | | |
| 50[247]48[240] | GTCACGCTGGGCGATGAGAATAAA | | | |
| 50[279]48[272] | GAGCGGGCAATCAAGTTACCGACC | | | |
| 50[311]48[304] | GTGGCGAGAGCACTAACATCTTCT | | | |
| 50[327]51[335] | GGGAAAGCCTGAGAAGTGTTTTT | | | |
| 50[375]48[368] | AGAGGCGAAACAAAATTGCAAATC | | | |
| 50[407]48[400] | TTGCTTTGTGAATTACGTTGGGTT | | | |
| 50[439]48[432] | CATCGGGAAAATCAATTGAGAGAC | | | |

| Oligo Name | Sequence (5' to 3') | Spring | Steric-1x | Steric-6x |
|---|---|---|---|---|
| 51[232]53[239] | AACGTGCAACTCAAGAACCCT | | | |
| 51[264]53[271] | AGCTAAACATTAGTAAGTGGCACA | | | |
| 51[296]53[303] | AGACAGGAGCAAATTAAGTCTTTA | | | |
| 51[336]50[328] | ATAATAAGTTTGAGTAACCTGAGCAAATAGAGCTTGACG | | | |
| 51[360]53[367] | ATTTTGCGCCTTTGCCAGCAGAAG | | | |
| 51[392]53[399] | GCGGAATTTTTACAAAGTATTAAC | | | |
| 51[424]53[431] | TGATGGCTAATACAAGAGCCA | | | |
| 52[247]50[248] | AGTAGAAGTTTCCTCGTTAGAATCGTGTAGCG | | | |
| 52[279]50[280] | CTTCTTTGAGGAGGCCGATTAAAGCGAAAG | | | |
| 52[311]50[312] | TCCATCACACGGTACGCCAGAATCCGGCGAAC | | | |
| 52[351]58[336] | ATTAATTTTAACAGTGAGGCCACCATCGCCATTAAAAACATTAAAG | | | |
| 52[375]50[376] | TATTAAATGAACAAAGAAACCACATCGCGC | | | |
| 52[407]50[408] | TATTAGACATCATCATATTCCTGATCGCCTGA | | | |
| 52[439]50[440] | TCAATAGAAATTCATCAATATAATACCTTTTA | | | |
| 53[224]52[216] | CAGAGATAACTATCGGCCTTGCTG | | | |
| 53[240]57[254] | TCTGACCTGAAACGTCATCAGTAGCGA TTTT | | | |
| 53[272]59[279] | GACAATATCACCAGTAGCACCATTCACCCTCAGAGCCGCC | | | |
| 53[304]59[311] | ATGCGCGATGAGCCATTTGGGAATAGAGCCGCCGCCAGCA | | | |
| 53[368]59[375] | ATAAAACAGGGAGGGAAGGTAAATAGCTATCTTACCGAAG | | | |
| 53[400]59[407] | ACCGCCTGAAAGACAAAAGGGCGAGTAAGCAGATAGCCGA | | | |
| 53[432]59[439] | GCAGCAAACAATAGAAAATTCATGAAACCGAGGAAACG | | | |
| 54[223]50[216] | TTCTGACGTTTGACGAACCCGCCG | | | |
| 54[248]59[239] | TTTT GACCAGTAATAACACCGTAACCAATGCCACCCT | | | |
| 56[219]49[231] | GTAGTTTACATTTACTAGAAAAAGGGCGAAAAACCG | | | |
| 56[254]62[240] | TTTT CAGAATCAAGTGCCAACATCCACCACC | | | |
| 57[208]61[215] | CCCCTTATACGGGGTCTTAATGCC | | | |
| 58[255]52[248] | CAAGGCCGGAAAGCGTAAGAATACTAACATCACTTGCCTG | | | |
| 58[287]52[280] | GCAAAATTTTTGAATGGCTATTACCGTTGTAGCAATA | | | |
| 58[319]52[312] | CACCGACTACTGATAGCCCTAAAACGAGTAAAAGAGTCTG | | | |
| 58[335]58[352] | GTGAATTGCAGGTCAGACGATTGGCATGAAATAGCAATATTGACGG | | | |
| 58[351]52[352] | AAATTATTTACCGAACGAACCACCCGAACGTT | | | |
| 58[383]52[376] | CGATTGAGAGGTGAGGCGGTCACAATTCGACAACTCG | | | |
| 58[415]52[408] | CCAGCGCCCAACAGTGCCACGCTGTTTGAGGATTTAGAAG | | | |
| 58[447]52[440] | GTCACAATTGAAAAATCTAAAGCATAATAGATTAGAGCCG | | | |
| 59[216]56[220] | GCCTCCCTAAGTTTTATAGCGTTTGACT | | | |
| 59[240]60[248] | CAGAACCGCCACCCTCGCTTTTGA | | | |
| 59[280]61[279] | ACCAGAAGAATTTACCGTTCCAACTCCTCA | | | |
| 59[312]61[311] | TTGACAGGTCATTAAAGCCAGAATGTTTTGCT | | | |
| 59[376]61[375] | CCCTTTTGAATTGAGTTAAGCCCCCAGCTA | | | |
| 59[408]61[407] | ACAAAGTTTTGAGCGCTAATATCAACGCTAAC | | | |
| 59[440]61[439] | CAATAATAACTGAACACCCTGAACTGCCAGTT | | | |
| 60[231]53[223] | CTGGTAATCAGAGCCGAAACCATCGCCATCTTAAAGCCAA | | | |
| 60[247]63[255] | TGATACAGATTCTGAAACATGAAACCTCAGAGGTAATTTA | | | |
| 60[263]58[256] | CATACATGAGAGCCACACCATTAG | | | |
| 60[295]58[288] | CAGTCTCTCCACCACCTAGAGCCA | | | |
| 60[327]58[320] | ATAAATCCAGGTTGAGATCACCGT | | | |

| Oligo Name | Sequence (5' to 3') | Spring | Steric-1x | Steric-6x |
|---|---|---|---|---|
| 60[359]61[351] | AGAGCAAGAAACACTTGATATTCATCGAGAGGGTTAAGAT | | | |
| 60[391]58[384] | ACCCACAATAAGAAAACATTCAAC | | | |
| 60[423]58[416] | GAGGGTAAACCAGAAGATGGTTTA | | | |
| 61[216]63[223] | CCCTGCCTAGCAAGCCTGAGAATC | | | |
| 61[228]54[224] | CGGATAGCGTCAGATAGCAGAAGGGACA | | | |
| 61[280]63[287] | AGAGAAGCCCTCAGATAAGAG | | | |
| 61[312]63[319] | CAGTACCATATCACCGAAGTAATT | | | |
| 61[352]60[360] | TAGTTGCTATTTTGCACAATAATA | | | |
| 61[376]63[383] | CAATTTTCCTCCCGTATCAAC | | | |
| 61[408]63[415] | GAGCGTCTGGTATTCTAATAATAT | | | |
| 61[440]63[447] | ACAAAATAAGCAAGCAAGAAACCA | | | |
| 62[239]61[227] | CTCATTTTCAGGGATATTT | | | |
| 62[263]60[264] | ACCGCCACGTATTAAGAGGCTGAGGTAAGCGT | | | |
| 62[295]60[296] | TACCGCCAGATTAGGATTAGCGGGGGAAAGCG | | | |
| 62[327]60[328] | AATAGGTGGGCGGATAAGTGCCGCAAACAA | | | |
| 62[359]63[351] | GAAGCCTTAAATCGATAACATGTT | | | |
| 62[391]60[392] | TTAGCGAAATCCTGAATCTTACCAGAGAGATA | | | |
| 62[423]60[424] | GCTTATCCTTCCAGAGCCTAATTAAAGTCA | | | |
| 63[224]51[231] | GCCATATTATCATAATGGCAGATTACCACCACGCACGTAT | | | |
| 63[256]49[263] | GGCAGAGGAATAAGGCGTTAAATAGCCCACTACGTGAACC | | | |
| 63[288]49[295] | AATATAAATTTAATGGTTTGAAATTTTTGGGGTCGAGG | | | |
| 63[320]49[327] | CTGTCCAGATATTTTAGTTAATTTATCGGAACCCTAAAGG | | | |
| 63[352]49[359] | CAGCTAATAGACAAAGAACGCGAGAGATGATGAAACAAAC | | | |
| 63[384]49[391] | AATAGATAATATGTAAATGCTGATAATTACATTTAACA | | | |
| 63[416]49[423] | CCCATCCTTTAACCTCCGGCTTAGCTTTTTTAATGGAAAC | | | |
| 48[463]62[456] | GTGAATTTTCGGCTGTCTTTCCTTGTAGGAATCATTACCG | | | |
| 49[456]51[455] | CCTTGCTTAATATACAGTAACAGTCCTGATTG | | | |
| 49[488]51[487] | ATTTTCCCTTGCGTAGATTTTCAGGGAAGGGT | | | |
| 50[471]48[464] | GTCAGATGCTGTAAATAGTCAATA | | | |
| 51[456]53[463] | TTTGGATTCTAACAACTCACCTTG | | | |
| 51[488]53[495] | TAGAACCTAGGAAGGTCAATCAAT | | | |
| 52[471]50[472] | TAGGAGCAATACTTCTGAATAATGTTTAAC | | | |
| 53[464]59[471] | CTGAACCTGACACCACGGAATAAGCCCAAAAGAACTGGCA | | | |
| 58[479]52[472] | ACGCAAACAAATATCAAACCCTTATCTAAAATATCTT | | | |
| 59[472]61[471] | TGATTAAAAACAGGGAAGCGCAATCCCAAT | | | |
| 60[455]58[448] | GAGAATTAACGGAATATTTATTTT | | | |
| 60[487]58[480] | TAACATAAGACTCCTTTAAAAGAA | | | |
| 61[472]63[479] | CCAAATATTTCATCATCATTC | | | |
| 62[455]60[456] | CGCCCAATAACAGCCATATTATTTTTAGACGG | | | |
| 63[448]49[455] | ATCAATAAATCAAAATCATAGGTCATATGTGAGTGAATAA | | | |
| 63[480]49[487] | CAAGAACGTAAGACGCTGAGAAGCGTCGCTATTAATTA | | | |
| 48[519]63[519] | GGCGCTAGGGCAAACATAGCGATAGCTTAGATGGTATTAAACCAAGTACCGCAAAACGCAATAA | | | |
| 50[527]49[513] | CCAGCAGAAGCGTAAAACAGAAATAAAGAAATTAGAATCCTTGAGCTGG | | | |
| 52[519]51[521] | TACAACAACAGTTGAAAGGAATTGACCATATCAAAATTATTTGCAATAAA | | | |
| 53[496]59[519] | ATCTGGTCTAAAGGTGGCAACATAATTACGCAGTATGTTAGCAAACGTAGAAACCT | | | |
| 58[519]53[519] | TGAATAAATACATACAAGTTGGCAAATAATCG | | | |

| Oligo Name | Sequence (5' to 3') | Spring | Steric-1x | Steric-6x |
|---|---|---|---|---|
| 60[519]61[519] | ACGAGCATAAATAGCAGCCTTTACTTAACGTCAAAAATGAGTAGAAAC | | | |
| 62[516]60[488] | AACCGAGGCTCATCGAGAACAAGCAAGCCGTTTTTATAGAAACGATTTTTTGTAGAGAGAA | | | |
| 5[15]2[9]_Biotin | TTTT AACGAGTAGTAAATGTACAGACCAGGCGCATA TTTT/3Biosg/ | | | |
| 9[15]6[9]_Biotin | TTTT CGAGGCATAGTAAGGACGTTGGGAAGAAAAAT TTTT/3Biosg/ | | | |
| 13[15]10[9]_Biotin | TTTT TACCCTGACTATTAGGGGGTAATAGTAAAATG TTTT/3Biosg/ | | | |
| 15[32]14[9]_Biotin | GATTAGAGAGCTTCAAAGCGAACCAGACCGGAA TTTT/3Biosg/ | | | |
| Biotin_19[12]18[12] | /5Biosg/TTTT TTGATTCCCAATTCTGTCATTCCATATAACAG TTTT | | | |
| Biotin_23[12]24[9] | /5Biosg/TTTT TCCAATAAACCAAAAACAT TTTT | | | |
| Biotin_25[9]26[15] | /5Biosg/TTTT TATGACCCTGTAATACTTTTATATTTTAAATG TTTT | | | |
| Biotin_29[9]30[15] | /5Biosg/TTTT ATGCCGGAGAGGGTAGCTAGTAATCGTAAAAC TTTT | | | |
| Blotin_33[9]34[15] | /5Biosg/TTTT AATTCGCATTAAATTTTTGCTTTCATCAACAT TTTT | | | |
| Biotin_37[9]38[15] | /5Biosg/TTTT TGCATCTGCCAGTTTGAGGACCAGGCAAAGCG TTTT | | | |
| 41[12]40[12]_Biotin | TTTT CGATTAAGTTGGGTAAGGGATGTGCTGCAAGG TTTT/3Biosg/ | | | |
| 45[12]44[12]_Biotin | TTTT TTGTGTACATCGACATGCCAGCAGTTGGGCGG TTTT/3Biosg/ | | | |
| 25[48]24[65] | TATTTCAACGCTATTGGGCGCCAGGCCTGGC | | | |
| 27[48]29[55] | TTCAAAAGGATAAAAACAAAGGCT | | | |
| 52[215]54[208] | GTAATATCATGGTTGCCTCATGGA | | | |
| 55[208]56[193] | ATGGATTACGCGTTTTCATCGGCATTT TTTT | | | |
| 63[203]62[189] | AGTAGGGCTTAATCAATAGGAACCCATGTACCGTAACACT | | | |
| 50[215]48[189] | CGCTTAATCCAACGTCAAAGCCTGTTTAGTATCATATGCGTTA | | | |
| 49[189]50[193] | CTATTAAAGAACGTGGACTGCGCCGCTACA | | | |
| 51[193]52[191] | GGGCGCGTACTCAGAACAATATTA | | | |
| 53[191]58[208] | CCGCCAGCCATTGCAACAGGATTCATAAT | | | |
| 55[188]54[188] | CGCTCAATCGTCTGAAAATACCTACATTTTGA | | | |
| 57[193]58[191] | TCGGTCATAGCCAAAATCACCGGA | | | |
| 59[191]59[215] | ACCAGAGCCACCACCGGAACC | | | |
| 61[188]60[188] | GTGCCCGTATAAACAGAGTGCCTTGAGTAACA | | | |
| 24[55]29[55] | AAGCCTCATTCAAAAGGATAAAAACAAAGGCT | | | |
| 25[48]24[56] | TATTTCAACGCTATTGGGCGCCAGGCCTGGCCCTGCAATA | | | |
| 50[215]48[189] | CGCTTAATCCAACGTCAAAGCCTGTTTAGTATCATATGCGTTA | | | |
| 63[208]62[189] | GGCTTAATCAATAGGAACCCATGTACCGTAACACT | | | |
| 52[215]56[208] | GTAATATCATGGTTGCCTCATGGAATGGATTACGCGTTTT | | | |

**Table S2. Elastic spring system staples.**

| Oligo Name | Sequence (5' to 3') | L1-21nt | DAapt | SRTNapt |
|---|---|---|---|---|
| **L1**_63[189]24[48] | AGCCAACGCTCAAC **CGTTGACCAGACTCATTGTCG** CCTGCAATAAAGCCTCA | | | |
| **DAapt**_63[189]24[48] | AGCCAACGCTCAAC **CGACGCCAGTTTGAAGGTTCGTTCGCAGGTGTGGAGTGACGTCG** CCTGCAATAAAGCCTCA | | | |
| **SRTNapt**_63[189]24[48] | AGCCAACGCTCAAC **CGACTGGTAGGCAGATAGGGGAAGCTGATTCGATGCGTGGGTCG** CCTGCAATAAAGCCTCA | | | |

### Table S3. Steric hindrance system staples

| Oligo Name | Sequence (5' to 3') | 1x-Proximal | 1x-Midpoint | 1x-Peripheral |
|---|---|---|---|---|
| 49[189]50[193] | CTATTAAAGAACGTGGACTGCGCCGCTACA | | | |
| 51[193]52[191] | GGGCGCGTACTCAGAACAATATTA | | | |
| 53[191]58[208] | CCGCCAGCCATTGCAACAGGATTCATAAT | | | |
| 55[188]54[188] | CGCTCAATCGTCTGAAAATACCTACATTTTGA | | | |
| 57[193]58[191] | TCGGTCATAGCCAAAATCACCGGA | | | |
| 59[191]59[215] | ACCAGAGCCACCACCGGAACC | | | |
| 61[188]60[188] | GTGCCCGTATAAACAGAGTGCCTTGAGTAACA | | | |
| 63[189]56[193] | AGCCAACGCTCAACAGTAGCATCGGCATTT | | | |
| 63[189]56[193]_**L1** | AGCCAACGCTCAACAGTAGCATCGGCATTT **CGTTGACCAGACTCATTGTCG** | | | |
| | DNA-PAINT variant (PAINT handle and ATTO655-labelled imager strands): | | | |
| 63[189]56[193]_**P1** | AGCCAACGCTCAACAGTAGCATCGGCATTT **ATACATCTA** | | | |
| **P1**-ATTO655 | TAGATGTAT/3ATTO655/ | | | |
| 55[188]54[188]_**L1** | CGCTCAATCGTCTGAAAATACCTACATTTTGA **CGTTGACCAGACTCATTGTCG** | | | |
| 57[193]58[191]_**L1** | TCGGTCATAGCCAAAATCACCGGA **CGTTGACCAGACTCATTGTCG** | | | |

| Oligo Name | Sequence (5' to 3') | 6x |
|---|---|---|
| 53[191]58[208] | TTTT CCGCCAGCCATTGCAACAGGATTCATAAT | |
| 59[191]59[215] | TTTT ACCAGAGCCACCACCGGAACC | |
| 61[188]60[188]_**L1** | GTGCCCGTATAAACAGAGTGCCTTGAGTAACA **CGTTGACCAGACTCATTGTCG** | |
| 63[189]56[193]_**L1** | AGCCAACGCTCAACAGTAGCATCGGCATTT **CGTTGACCAGACTCATTGTCG** | |
| 57[193]58[191]_**L1** | TCGGTCATAGCCAAAATCACCGGA **CGTTGACCAGACTCATTGTCG** | |
| 55[188]54[188]_**L1** | CGCTCAATCGTCTGAAAATACCTACATTTTGA **CGTTGACCAGACTCATTGTCG** | |
| 49[189]50[193]_**L1** | CTATTAAAGAACGTGGACTGCGCCGCTACA **CGTTGACCAGACTCATTGTCG** | |
| 51[193]52[191]_**L1** | GGGCGCGTACTCAGAACAATATTA **CGTTGACCAGACTCATTGTCG** | |

### Table S4. Lever arm staples.

| Oligo Name | Sequence (5' to 3') |
|---|---|
| 48[559]62[552] | AACGTGGCAAGACTCCTTATTACGGCCCTTTTTAAGAAAA |
| 48[591]62[584] | CCGATTTACGTAGAAAATACATACACAATGAAATAGCAAT |
| 48[623]62[616] | AAGCACTATAAAAGAAACGCAAATTGAGTTAAGCCCA |
| 48[655]62[648] | CAAATCAAGTTTATTTTGTCACAAGAGCGCTAATATCAGA |
| 48[687]62[680] | AGGGCGATTATGGTTTACCAGCGCTGAACACCCTGAACAA |
| 48[719]62[712] | TCCAACGACATTCAACCGATTGACAGGGAAGCGCATT |
| 48[751]62[744] | GGAACAAGTATTGACGGAAATTATATAGCAGCCTTTACAG |
| 48[783]62[776] | CCCGAGATTATCACCGTCACCGACAACGATTTTTTGTTTA |
| 48[815]62[808] | CGGCAAATTAGAGCCAGCAAAAAGCCATATTATTTAT |
| 48[847]62[847] | GCGAAAATTACCATTAGCAAGGCCTGTATCACC |
| 48[879]62[872] | GCAAGCGGGAAACCATCGATAGCAGAGGGTTGATATAAGT |
| 48[911]62[904] | CCCTTCACGACAGAATCAAGTTTGCTCAGTACCAGGC |
| 48[943]62[936] | TTTTCACCCTGTAGCGCGTTTTCACCTCAAGAGAAGGATT |
| 48[975]62[968] | GTTTGCGTATAGCCCCCTTATTAGTGAAACATGAAAGTAT |
| 48[1007]62[1000] | TAATGAAATAATCAAAATCACCTGCCCCCTGCCTATT |
| 48[1039]62[1032] | TCCAGTCGCCGGAACCGCCTCCCTCCTTGAGTAACAGTGC |
| 48[1071]62[1064] | ACATTAATCAGAACCGCCACCCTCGTACTGGTAATAAGTT |
| 48[1103]62[1096] | AGCCTGGGAGCCGCCACCAGAACGTCATACATGGCTT |
| 49[514]48[520] | CAAGTGTAGCGGTCACGCTGCAAAGCGAAAGGAGCG |
| 49[552]51[551] | CGCCGCGCAAACATCGCCATTAAATTAACACC |
| 49[584]51[583] | TACTATGGTTAGTCTTTAATGCGCAGCCAGCA |
| 49[616]51[615] | TGCTTTCACGTGGCACAGACAACCTTGCTG |
| 49[648]51[647] | AAACAGGAATAGAACCCTTCTGACTCAATATC |
| 49[680]51[679] | AGGAACGGAGTAATAAAAGGGACATGAAAGGA |
| 49[712]51[711] | TTTATAATTACATTGGCAGATTCTTTAGGA |
| 49[744]51[743] | GTCTGTCCATTTTGACGCTCAATCGCCGTCAA |
| 49[776]51[775] | CAATACTTCAACAGGAAAAACGCTGAAGTATT |
| 49[808]51[807] | GCCTGAGATATCCAGAACAATATCGTATTA |
| 49[840]50[824] | ATGAGCCGCAATCCGCCGGGCGCGATCGGCCT |
| 49[872]51[871] | TAATGGGTTCAGCGGGGTCATTGCTCTTCGCT |
| 49[904]51[903] | CATCCCTGCAACCAGCTTACGGATGTGCTG |
| 49[936]51[935] | TTAACGGCCAACGGCAGCACCGTCCCAGGGTT |
| 49[968]51[967] | CCAGCGGTCTGGTCTGGTCAGCAGCGACGGCC |
| 49[1000]51[999] | TCCAGCGCGTGCCGGACTTGTACCGCCACG |
| 49[1032]51[1031] | TCTGTGGTGCCTCCGGCCAGAGCACAATCGGC |
| 49[1064]51[1063] | CGTTTTCACGCGGTCCGTTTTTTCCAGTCCCG |
| 49[1096]51[1095] | GCACGCGGTTAAACGATGCTGACCGTGGTG |
| 50[567]48[560] | TAGCCCTATTAATGCGAGCCGGCG |
| 50[599]48[592] | AATGGCTATTGCTTTGGGGAGCCC |
| 50[631]48[624] | GTAAGAATCTCGTTAGGTGCCGTA |
| 50[663]48[656] | CAACAGAGGGCCGATTCCATCACC |
| 50[695]48[688] | ACACGACCTACGCCAGCGTCTATC |
| 50[727]48[720] | TGGATTATTCAGTGAGACGTGGAC |
| 50[759]48[752] | ATACCTACATCACGCATCCAGTTT |
| 50[791]48[784] | AGCCATTGCTTTGATTAAGAATAG |
| 50[823]48[816] | TGCTGGTATAGAAGAATCCGAAAT |

| Oligo Name | Sequence (5' to 3') |
|---|---|
| 50[855]48[848] | TTCGCACTGGTCACTGCCCAGCAG |
| 50[887]48[880] | GTCCAGCAAAAGGTTTGAGTTGCA |
| 50[919]48[912] | CATCCCACTACACTGGGCTGATTG |
| 50[951]48[944] | AAGAATGCATCAGATGGGTTTTTC |
| 50[983]48[976] | GCGTGGTGGCCGGTGCGAGAGGCG |
| 50[1015]48[1008] | TAACGGAACAGTGTCAAGCTGCAT |
| 50[1047]48[1040] | CGCTGGCAGCTGCGGCGCCCGCTT |
| 50[1079]48[1072] | CCGGCAAACGGTCATAGCTAACTC |
| 50[1111]48[1104] | AGGGTAAATGCCTGTTAAGTGTAA |
| 51[522]55[535] | ACAGAGGTGAGGCAAGTACGGCAATGAAT |
| 51[552]53[559] | GCCTGCAAAACAATAAACAGTTGA |
| 51[584]53[591] | GCAAATGAATACAGTAATTTAGTT |
| 51[616]53[623] | AACCTCACGTAGATTGGTCAA |
| 51[648]53[655] | TGGTCAGTTATTTGCATTGGGGCG |
| 51[680]53[687] | ATTGAGGAATGGAAGGTCTACTAA |
| 51[712]53[719] | GCACTAAATCCTGAAATCATA |
| 51[744]53[751] | TAGATAATTGATTATCATTAAGCA |
| 51[776]53[783] | AGACTTTAACCAGAAGAAATCGGT |
| 51[808]53[815] | AATCCTTTAACATTTAATACT |
| 51[840]53[847] | GGGAAGGGCGCCATTCACGCAAGG |
| 51[872]53[879] | ATTACGCCTCTGGTGCATATTTTA |
| 51[904]53[911] | CAAGGCGATCGCACGGTAAAG |
| 51[936]53[943] | TTCCCAGTGAGGGGACAGACAGTC |
| 51[968]53[975] | AGTGCCAAGGGCGCATAACCGTTC |
| 51[1000]53[1007] | GGAACGGCCGTAATGGGTAGC |
| 51[1032]53[1039] | GAAACGTACGGATTCTTATCAGGT |
| 51[1064]53[1071] | GAATTTGTCAACATTAACAAGAGA |
| 51[1096]53[1103] | AAGGGATCGCGTCTTAGCATG |
| 52[543]53[535] | CCTGATTGCTTTGAATACCAAGTCGCAGAGTGTCTGGA |
| 52[567]50[568] | TCGGGAGACAGTGCCACGCTGAGGAACTGA |
| 52[599]50[600] | AGATGAATAAAATCTAAAGCATCATATTTTTG |
| 52[631]50[632] | AGAAATTGAATATCAAACCCTCAACTGAAAGC |
| 52[663]50[664] | ATCAAAATTGGCAAATCAACAGTTTCTGGC |
| 52[695]50[696] | TCTGAATAAGGTTATCTAAAATATCACCAGTC |
| 52[727]50[728] | TCAATATACAACTAATAGATTAGAGTCTGAAA |
| 52[759]50[760] | CATATTCCACATTTGAGGATTTACATGGAA |
| 52[791]50[792] | AAGAAACCCAAACAATTCGACAACTTACCGCC/3ATTO488N/ |
| 52[823]51[839] | AGTTTGAGTGCCCGAACGTTATTACAACTGTT |
| 52[855]50[856] | GCGCCATTCGATCGGTGCGGGCCAGGCGCT |
| 52[887]50[888] | GCACCGCTAGCTGGCGAAAGGGGGCTGGAGGT |
| 52[919]50[920] | TCAGGAAGATTAAGTTGGGTAACGGGTGGTGC |
| 52[951]50[952] | GCCAGTTTCACGACGTTGTAAAACAACCGC |
| 52[983]50[984] | GTGTAGATGCTTTCAGAGGTGGAGGAACGTCA |
| 52[1015]50[1016] | CGGATTGAATAACCTCACCGGAAACATCCTCA |
| 52[1047]50[1048] | CAACCCGTCAGCGCCATGTTTACGTCTCGT |
| 52[1079]50[1080] | GCTTTCATGAGAGATAGACTTTCTTTGCCGTT |

| Oligo Name | Sequence (5' to 3') |
|---|---|
| 53[536]50[528] | AGTTTCATTGCAACTAGGTCAGTAAATACCGAACGAACCA |
| 53[560]59[567] | TTCCCAATTTAGAGAGTACCTTTATTCCCTTAGAATCCTT |
| 53[592]59[599] | TGACCATTCAGACCGGAAGCAAACGCTTAGATTAAGACGC |
| 53[624]59[631] | TAACCTGTATCGCGTTTTAATTCTGAATTTATCAAAAT |
| 53[656]59[663] | CGAGCTGAAAGATTAAGAGGAAGCTACCTTTTTAACCTCC |
| 53[688]59[695] | TAGTAGTATATAGTCAGAAGCAAAATATAACTATATGTAA |
| 53[720]59[727] | CAGGCAAGAAATCAAAAATCAGGAATCGCAAGACAAAG |
| 53[752]59[759] | ATAAAGCCCTTTAAACAGTTCAGATTCAAATATATTTTAG |
| 53[784]59[791] | TGTACCAATCATAAATATTCATTGGACCTAAATTTAATGG |
| 53[816]59[823] | TTTGCGGGAGACTGGATAGCGTCTGTGATAAATAAGGC |
| 53[848]59[855] | ATAAAAATGTTTTGCCAGAGGGGGAACACTCATCTTTGAC |
| 53[880]59[887] | AATGCAATACCAAAATAGCGAGAGCAAGCGCGAAACAAAG |
| 53[912]59[919] | ATTCAAAACATAACCCTCGTTTAATCATCGCCTGATAA |
| 53[944]59[951] | AAATCACCGGAATTACGAGGCATACGACCTGCTCCATGTT |
| 53[976]59[983] | TAGCTGATATTCAACTAATGCAGAGGCGCAGACGGTCAAT |
| 53[1008]59[1015] | TATTTTTGAGATTCATCAGTTGATGACCAACTTTGAAA |
| 53[1040]59[1047] | CATTGCCTGAACTAACGGAACAACTGTACAGACCAGGCGC |
| 53[1072]59[1079] | ATCGATGACGTTGGGAAGAAAAATTTCATCAAGAGTAATC |
| 53[1104]59[1111] | TCAATCATTTTAAGAACTGGCTCATTCATTACCCAAAT |
| 54[558]52[544] | TTTT TGTTTTAAATATCCATATACGGATTCG |
| 55[497]54[520] | TTTT GCAAAAGAAGATGATGAAACAAACATGCGAATT |
| 55[536]49[551] | ATAATGCTAAGGGAAGGCGTAACCACCACACC |
| 56[519]54[496] | TTAATTACATGGAAACATTCATTTCAATTACCTGA TTTT |
| 56[552]60[520] | TTTT GGCTTAGAGCTTAATTGCGAAAACAAAACAATCAACTAATTT |
| 57[492]56[492] | TTTT TGAATTACCTTTTTTAATTTAACAATTTCATT TTTT |
| 57[520]61[535] | AGTACATAAATCGGTATCCCATCTAATCGGC |
| 58[543]59[535] | TTTTGATAAGAAATATATGTGAGTGCTTCTGTAAATCG |
| 58[575]52[568] | GTCAGGATCTGCGAACGAGTAGACAGTACCTTTTACA |
| 58[607]52[600] | AAGCGAACAGATACATTTCGCAAATTTCAGGTTTAACGTC |
| 58[639]52[632] | CTTCAAATTTAGCTATATTTTCATCGTAAAACAGAAATAA |
| 58[671]52[664] | CATCAAAAAAGGTGGCATCAATGTTAGAACCTACCAT |
| 58[703]52[696] | CTGACTATGCATTAACATCCAATATTGTTTGGATTATACT |
| 58[735]52[728] | ATGACCATGCAAAGAATTAGCAAAAGATGATGGCAATTCA |
| 58[767]52[760] | TCAAATGTCAGAGCATAAAGCTGAGCGGAATTATCAT |
| 58[799]52[792] | CGGAATCGAAACATTATGACCCTGATCATTTTGCGGAACA |
| 58[831]52[824] | AAATGTTTAGAAGCCTTTATTTCAAGGCTGCGATTTTAAA |
| 58[863]52[856] | AAAAGAATTTTAGAACCCTCATCGGAAACCAGGCAAA |
| 58[895]52[888] | CGATAAAAGCCTGAGTAATGTGTATCCAGCCAGCTTTCCG |
| 58[927]52[920] | AACACTATGGGTGAGAAAGGCCGGGACGACAGTATCGGCC |
| 58[959]52[952] | GCCAAAAATCAATATGATATTCCGTAACCGTGCATCT |
| 58[991]52[984] | AATACCACAAATTAATGCCGGAGAGGGATAGGTCACGTTG |
| 58[1023]52[1016] | AGGTAGAAAGAGATCTACAAAGGCCCGTGGGAACAAACGG |
| 58[1055]52[1048] | ATAAAACGAGAGTCTGGAGCAAAATGTGAGCGAGTAA |
| 58[1087]52[1080] | AGTCAGGAACGGTAATCGTAAAACGGCCTTCCTGTAGCCA |
| 59[536]57[552] | TCGCTATTAAAATAATCATTTTTGCGGAT TTTT |
| 59[568]61[567] | GAAAACATTTATCAACAATAGACGGGTATT |

| Oligo Name | Sequence (5' to 3') |
|---|---|
| 59[600]61[599] | TGAGAAGAACAACATGTTCAGCTAGAACAAGC |
| 59[632]61[631] | CATAGGTCTAAAGTAATTCTGTCCGGAATCAT |
| 59[664]61[663] | GGCTTAGTAATAAGAGAATATACAGATATA |
| 59[696]61[695] | ATGCTGATATGTAATTTAGGCAGAAACGCGAG |
| 59[728]61[727] | AACGCGAGATTGAGAATCGCCATATGCGGGAG |
| 59[760]61[759] | TTAATTTGTATAAAGCCAACGCAGTTGCTA |
| 59[792]61[791] | TTTGAAATTAGTATCATATGCGTTTCCTGAAT |
| 59[824]58[832] | GTTAAATAAGAATTAATAATAGTA |
| 59[856]61[855] | CCCCAGCAAAACGAAAGAGGCAGCCACCCT |
| 59[888]61[887] | TACAACGGAATACGTAATGCCACTCCACCCTC |
| 59[920]61[919] | ATTGTGTCTTTCATGAGGAAGTTTGGATAGCA |
| 59[952]61[951] | ACTTAGCCGGCTACAGAGGCTTTAACACTG |
| 59[984]61[983] | CATAAGGGAGCGAAAGACAGCATCCAACGCCT |
| 59[1016]61[1015] | GAGGACAGAGGCCGCTTTTGCGGGATAGTTAG |
| 59[1048]61[1047] | ATAGGCTTATTCGGTCGCTGAGCTTTCCAG |
| 59[1080]61[1079] | TTGACAAGTGACAACAACCATCGCATGGGATT |
| 60[551]58[544] | GAACAAGAAATTAATTATTGCTCC |
| 60[583]58[576] | CGCGCCTGTAGCGATATCCAACAG |
| 60[615]58[608] | GACAATAAGTCAATAGGAGCTTCA |
| 60[647]58[640] | ACAAAAGGTGAGAGACCCGAAAGA |
| 60[679]58[672] | CGAGCCAGGTTGGGTTGCGGATTG |
| 60[711]58[704] | ACGCCAACGCAAATCCTCTTTACC |
| 60[743]58[736] | AGGGCTTAAAAACTTTAAACGAGA |
| 60[775]58[768] | TCTTACCACATCTTCTAATCCCCC |
| 60[807]58[800] | AGCCTGTTACCGACCGCAATACTG |
| 60[831]61[823] | CCGGAATCATAATTACTTCCAGAG |
| 60[871]58[864] | ACCAACCTGATTATACGCTTTTGC |
| 60[903]58[896] | ACGGGTAAAGATTTGTCCAGACGA |
| 60[935]58[928] | AAAGACTTGAAATCCGGTAAGAGC |
| 60[967]58[960] | GGTAGCAACGGAACGATACATAAC |
| 60[999]58[992] | CCCTCAGCAACCGAACGATTTAGG |
| 60[1031]58[1024] | GGAGTTAAATGAACGGATTATTAC |
| 60[1063]58[1056] | AACCGATAGGCTGACCCTACGTTA |
| 60[1095]58[1088] | GCCGACAAAACCGGATATTATACC |
| 61[536]63[543] | TGTCTTTCATAGCCGAAAGAACTG |
| 61[568]63[575] | AAACCAATACCGAACAGTATG |
| 61[600]63[607] | AAGCCGTTAGCAAGAAATAAAGGT |
| 61[632]63[639] | TACCGCGCCCACAAGAAGACACCA |
| 61[664]63[671] | GAAGGCTGGTAATTTCAATAG |
| 61[696]63[703] | GCGTTTTAGAATTAACCAAAGACA |
| 61[728]63[735] | GTTTTGAAACATAAAAAGGGAGGG |
| 61[760]63[767] | TTTTGCAAATGAAATCATTAA |
| 61[792]63[799] | CTTACCAAAAATAAGATTGAGCCA |
| 61[824]60[832] | CCTAATTTGCCAGGGTTTAGTACCAAAGAATACACAAACA |
| 61[856]63[863] | CAGAACCGAATAGGGGAAACG |
| 61[888]63[895] | AGAGCCACGCCGTCGAGCACCGTA |

| Oligo Name | Sequence (5' to 3') |
|---|---|
| 61[920]63[927] | AGCCCAATCGGGGTTTTGCCTTTA |
| 61[952]63[959] | AGTTTCGTGAGACTTCGGCAT |
| 61[984]63[991] | GTAGCATTTATTATTCCGTTTGCC |
| 61[1016]63[1023] | CGTAACGAACAGTTAAGGAACCAG |
| 61[1048]63[1055] | ACGTTAGGTCAGTGCAGAGCC |
| 61[1080]63[1087] | TTGCTAAAACAGGAGTAGAGCCAC |
| 62[551]60[552] | GTAAGCAGCTTATCATTCCAAGAATAAGTCCT |
| 62[583]60[584] | AGCTATCTGTACCGCACTCATCGAATGCAGAA |
| 62[615]60[616] | ATAATAAGTTTATTTTCATCGTAAGACGAC |
| 62[647]60[648] | GAGATAACCCAATAGCAAGCAAATAAGTACCG |
| 62[679]60[680] | AGTCAGAGTATCCGGTATTCTAAGGGCATTTT |
| 62[711]60[712] | AGACGGGAGCGAACCTCCCGACTTTTAACA |
| 62[743]60[744] | AGAGAATAGCCTTAAATCAAGATTTCAACAGT |
| 62[775]60[776] | ACGTCAAACCCAGCTACAATTTTAATACAAAT |
| 62[807]60[808] | CCCAATCCCGCTAACGAGCGTCTTAGAAAA |
| 62[846]63[831] | GTACTCAGGATTACAAAATAAACTCACCAGT |
| 62[871]60[872] | ATAGCCCGGCCACCCTCAGAACCGACGAAGGC |
| 62[903]60[904] | GGATAAGTCACCCTCATTTTCAGCCATTAA |
| 62[935]60[936] | AGGATTAGAGGAACCCATGTACCGTGAGGACT |
| 62[967]60[968] | TAAGAGGCTCACCAGTACAAACTAGGAACGAG |
| 62[999]60[1000] | TCGGAACCCCACAGACAGCCCTCATCGTCA |
| 62[1031]60[1032] | CCGTATAATCTAAAGTTTTGTCGTGCTTGCAG |
| 62[1063]60[1064] | TTAACGGGTAAATGAATTTTCTGTCCACGCAT |
| 62[1095]60[1096] | TTGATGATCAACTTTCAACAGTTTAGTTGC |
| 63[520]62[517] | TAACGGAATACCCAAACAAAGTTACCAGAAGGA |
| 63[544]55[558] | GCATGATTGAGAAAGGGTAGCTCAACA TTTT |
| 63[576]49[583] | TTAGCAAAGAGCTTGACGGGGAACCGCTACAGGGCGCG |
| 63[608]49[615] | GGCAACATAAATCGGAACCCTAAAACGAGCACGTATAACG |
| 63[640]49[647] | CGGAATAAGTTTTTTGGGGTCGAGAATCAGAGCGGGAGCT |
| 63[672]49[679] | AAAATTCAGGCCCACTACGTGAAAAAGGGATTTTAGAC |
| 63[704]49[711] | AAAGGGCGTCAAAGGGCGAAAAACAATCCTGAGAAGTGTT |
| 63[736]49[743] | AAGGTAAAAGTCCACTATTAAAGAGCCACCGAGTAAAAGA |
| 63[768]49[775] | AGGTGAATAGGGTTGAGTGTTGTAATTAACCGTTGTAG |
| 63[800]49[807] | TTTGGGAAATCCCTTATAAATCAAAGTAATAACATCACTT |
| 63[832]49[839] | AGCACCATCCTGTTTGATGGTGGTCTCAAACTGTTGCGGT |
| 63[864]49[871] | TCACCAATTCCACGCTGGTTTGCTTGCCCTGCGGCTGG |
| 63[896]49[903] | ATCAGTAGCCGCCTGGCCCTGAGACTTTGCTCGTCATAAA |
| 63[928]49[935] | GCGTCAGAAGTGAGACGGGCAACATGTGTTCAGCAAATCG |
| 63[960]49[967] | TTTCGGTCATTGGGCGCCAGGGTCCGGGTTACCTGCAG |
| 63[992]49[999] | ATCTTTTCTCGGCCAACGCGCGGGCCCCTGCATCAGACGA |
| 63[1024]49[1031] | AGCCACCAGGAAACCTGTCGTGCCCTGCGCGCCTGTGCAC |
| 63[1056]49[1063] | GCCACCCTTGCGTTGCGCTCACTCAGAATGCGGCGGGC |
| 63[1088]49[1095] | CACCCTCAGGTGCCTAATGAGTGACCGGGGGTTTCTGCCA |
| M1_48[1135]62[1128] | taagtgagacccgtacatat TT CACAACATCGCCAGCATTGACAGGAGCGCAGTCTCTGAAT TT taagtgagacccgtacatat |
| M1_48[1184]63[1184] | taagtgagacccgtacatat TTTT CATAGCTGTTTCCTGTGTGAAAGACGATTGGCCTTGATATTC TTTT taagtgagacccgtacatat |
| M1_49[1128]51[1127] | taagtgagacccgtacatat TT CTCCTCACCGCACAGGCGGCCTTTGAGACGCA TT taagtgagacccgtacatat |

| Oligo Name | Sequence (5' to 3') |
|---|---|
| M1_49[1160]51[1159] | taagtgagacccgtacatat TT GCTCGAATCATCGACATAAAAAAATTCTGCTC TT taagtgagacccgtacatat |
| M1_50[1143]48[1136] | taagtgagacccgtacatat TT AAAAAAGCAGTTGAGGCAATTCCA TT taagtgagacccgtacatat |
| M1_50[1182]49[1184] | taagtgagacccgtacatat TTTT GCGGTTGTGTATCGTAATCATGGT TTTT taagtgagacccgtacatat |
| M1_51[1128]53[1135] | taagtgagacccgtacatat TT GAAACAGCTTTAACCAAATCAGAA TT taagtgagacccgtacatat |
| M1_51[1160]53[1190] | taagtgagacccgtacatat TT ATTTGCCGCGCATTAATATAAGCAAATATTTAAATTGTAAACG TTTT taagtgagacccgtacatat |
| M1_52[1111]50[1112] | taagtgagacccgtacatat TT AAATAATTAGCTCTCACGGAAAAAAGTGATGA TT taagtgagacccgtacatat |
| M1_52[1143]50[1144] | taagtgagacccgtacatat TT GCTCATTTGGATCAAACTTAAATTCCCGTA TT taagtgagacccgtacatat |
| M1_52[1190]51[1182] | taagtgagacccgtacatat TTTT TTAATATTTTGTTAAAATTCCAGCAGTTGG TTTT taagtgagacccgtacatat |
| M1_53[1136]59[1143] | taagtgagacccgtacatat TT AAGCCCCAAACTTTAATCATTGTGCTCATTCAGTGAATAA TT taagtgagacccgtacatat |
| M1_58[1119]52[1112] | taagtgagacccgtacatat TT TATGCGATATGTACCCCGGTTGATATAGGAACGCCATCAA TT taagtgagacccgtacatat |
| M1_58[1151]52[1144] | taagtgagacccgtacatat TT TAATTTCAAAACAGGAAGATTGATTTTTGTTAAATCA TT taagtgagacccgtacatat |
| M1_58[1179]59[1179] | taagtgagacccgtacatat TTTT TAGTAAATTGGGCTTGGAAACACCAGAACGAG TTTT taagtgagacccgtacatat |
| M1_59[1112]61[1111] | taagtgagacccgtacatat TT CAACGTAAAACAGCTTGATACCGATCAGCGGA TT taagtgagacccgtacatat |
| M1_59[1144]61[1143] | taagtgagacccgtacatat TT GGCTTGCTTATCAGCTTGCTTTAGGAATTG TT taagtgagacccgtacatat |
| M1_60[1127]58[1120] | taagtgagacccgtacatat TT ATTTCTTACAAAGCTGAATTACCT TT taagtgagacccgtacatat |
| M1_60[1159]58[1152] | taagtgagacccgtacatat TT GTATCGGTCCTGACGAAGATGGTT TT taagtgagacccgtacatat |
| M1_60[1187]61[1187] | taagtgagacccgtacatat TTTT AGGCTCCAAAAGGAGCAAAATCTCCAAAAAAA TTTT taagtgagacccgtacatat |
| M1_61[1112]63[1119] | taagtgagacccgtacatat TT GTGAGAATCCAGTAAGCCACCACC TT taagtgagacccgtacatat |
| M1_61[1144]63[1151] | taagtgagacccgtacatat TT CGAATAAAATGGAAAGGTTGA TT taagtgagacccgtacatat |
| M1_62[1127]60[1128] | taagtgagacccgtacatat TT TTACCGTTAGAAAGGAACAACTAACGAGGTGA TT taagtgagacccgtacatat |
| M1_62[1159]60[1160] | taagtgagacccgtacatat TT AAAGCCAGTAATTTTTTCACGTTGCTTTAATT TT taagtgagacccgtacatat |
| M1_62[1184]62[1160] | taagtgagacccgtacatat TTTT ACAAACAAATAAATCCTCATT TT taagtgagacccgtacatat |
| M1_63[1120]49[1127] | taagtgagacccgtacatat TT AGAGCCGCACGAGCCGGAAGCATACTTCGCGTCCGTGAGC TT taagtgagacccgtacatat |
| M1_63[1152]49[1159] | taagtgagacccgtacatat TT GGCAGGTCATTGTTATCCGCTCAATCCCCGGGTACCGA TT taagtgagacccgtacatat |
| ATTO655-M1-5'-LNA | /5ATTO655/ttatatgtacgg |
| ATTO655-M1-3'-LNA | gtctcacttatt/3ATTO655/ |
